# Identifying hierarchical cell states and gene signatures with deep exponential families for single-cell transcriptomics

**DOI:** 10.1101/2022.10.15.512383

**Authors:** Pedro F. Ferreira, Jack Kuipers, Niko Beerenwinkel

## Abstract

Single-cell gene expression data characterizes the complex heterogeneity of living systems. Tissues are composed of various cells with diverse cell states driven by different sets of genes. Cell states are often related in a hierarchical fashion, for example, in cell differentiation hierarchies. Clustering which respects a hierarchy, therefore, can improve functional interpretation and be leveraged to remove noise and batch effects when inferring gene signatures. For this task, we present single-cell Deep Exponential Families (scDEF), a multi-level Bayesian matrix factorization model for single-cell RNA-sequencing data. The model can identify hierarchies of cell states and be used for dimension reduction, gene signature identification, and batch integration. Additionally, it can be guided by known gene sets to jointly type cells and identify their hierarchical structure, or to find higher resolution states within the provided ones. In simulated and real data, scDEF outperforms alternative methods in finding cell populations across biologically distinct batches. We show that scDEF recovers cell type hierarchies in a whole adult animal, identifies a signature of response to interferon stimulation in peripheral blood mononuclear cells, and finds both patient-specific and shared cell states across nine high-grade serous ovarian cancer patients.

## Introduction

Single-cell RNA-sequencing (scRNA-seq) data generates snapshots of cell states in a high-throughput fashion by quantifying expression levels of all genes, for thousands of individual cells [1]. These measurements permit the discovery of different cell populations within a tissue and the specific genes that are activated in each population. By aggregating scRNA-seq data from multiple donors these data can reveal commonalities in cell subpopulations between different individuals [2]. Analyzing tumor samples in this fashion has the potential to inform targeted therapy of cancer patients.

In recent years, a large number of methods to analyse scRNA-seq data have been developed, including for dimensionality reduction [3], clustering [4], differential expression analysis [5], pseudo-time analysis [6] and trajectory inference [7]. The analysis of scRNA-seq data typically involves the following steps: normalization, feature selection, dimensionality reduction, clustering, marker identification, and cluster annotation. Often, these steps rely on different computational methods or statistical models, each with their own assumptions. For example, the workflow described in [8] recommends dimensionality reduction with PCA on log-normalized UMI counts, followed by graph-based clustering, and differential expression between the clusters to identify the genes that define each cluster using a Wilcoxon rank-sum test. However, this multi-step approach is prone to inconsistencies in modelling assumptions and to statistical issues, such as artificially low p-values in differential gene expression [9].

Other approaches, like scHPF [10], simultaneously perform dimensionality reduction, clustering, and marker identification on the unnormalized UMI counts via a single probabilistic matrix factorization model, in which each non-negative factor describes a gene program. By relying on a model of the UMI counts, this method is able to produce gene signatures and cell groups jointly. However, it requires post-processing steps to choose the appropriate number of factors and does not distinguish between biological and technical variation in them. Another factorization-based method, f-scLVM [11] is based on the assumption that technical variation is not sparse and affects many genes simultaneously, but it does not enforce non-negativity.

None of these approaches explicitly models the inherent hierarchy of cell states. The hierarchical structure arises as some cell types may be further divided into subtypes (for example, regulatory or helper T cells) or into their specific functional roles (for example, cancer cells promoting angiogenesis or undergoing an epithelial-mesenchymal transition) [12]. The hierarchy reflects the specialization level at which cells perform their function, and it is therefore a natural feature to include in gene signature identification methods. Indeed, learning cell states at varying levels of resolution has been identified as one of the main challenges in single-cell data science [13]. Recently, nSBM [14] has been proposed to identify hierarchies of cell states by learning different levels at which cells can be clustered. However, by working on a k-nearest neighbor graph instead of the gene expression matrix, the method does not provide the gene signatures that define those clusters. Similarly, scHPL [15] learns trees of cell states but requires cell annotations, which may not be available.

An additional challenge in scRNA-seq data analysis is the integration of data from different sources, such as human samples from different donors, for example. Even when the samples contain functionally similar cell populations, technical issues, including batch effects, hide those similarities [16]. Two main approaches exist to account for batch effects. One approach is late integration, in which cells from different samples are first projected onto low-dimensional spaces and annotated separately, and then cells with the same labels across data sets (so-called anchor cells) are used to align the embeddings [17, 18, 19]. Another approach is to integrate the data earlier by appending data sets and using indicator variables in a statistical model to remove the sample-specific signal. This is done, for example, in ZINB-WaVE [20] and the neural network-based approach scVI [21].

The main challenge in data integration is the ability to remove sample-specific technical noise while retaining the true underlying biological variability [22]. On one hand, unsupervised matrix factorization models such as PCA, non-negative matrix factorization (NMF) or scHPF are prone to capturing technical batch effects. On the other hand, methods which use batch annotations to maximize the similarity between batches, such as scVI or Harmony [17], may lead to over-correction and the loss of true batch-specific cell states. We posit that using a hierarchy to impose similarities between factors contributes to minimizing technical batch effects while keeping true batch-specific signal.

Here, we present scDEF (single-cell Deep Exponential Families), a Bayesian model designed to identify hierarchical structure in scRNA-seq data. scDEF is a deep exponential family model [23] tailored to single-cell data in order to describe cells at multiple levels of resolution, which are directly associated with different gene signatures. By enforcing non-negativity, promoting sparsity, including hierarchical relationships among factors, and performing automatic model selection, scDEF is a general tool for hierarchical gene signature identification in scRNA-seq data for both single- and multiple-batch scenarios.

In the following sections, we first provide an overview of the scDEF model. On a benchmark study, we obtain improvements in performance over alternative methods, especially when different batches contain different cell populations. We demonstrate the basic analyses permitted by scDEF on peripheral blood mononuclear cells (PBMCs), and recover accurate cell type hierarchies in a whole adult animal. We show that our method performs robust batch integration and detects signatures of gene expression related to interferon stimulation in PBMCs. Finally, we apply scDEF to a cohort of nine high-grade serous ovarian cancer (HGSOC) patients and find cell states which are common across patients, and also patient-specific cell populations.

## Results

### Model overview

scDEF models gene expression heterogeneity among cells of a tissue as a set of sparse factors containing gene signatures for different cell states. These factors are related to each other through higher-level factors that contain more coarse-grained signatures. The method takes as input scRNA-seq count data and employs variational inference to learn a multi-level matrix factorization that describes the gene programs which are active in the data at multiple levels of resolution, depending on the number of layers (Figure 1 A). scDEF is based on DEFs [23] adapted to single-cell data.

**Figure 1:**
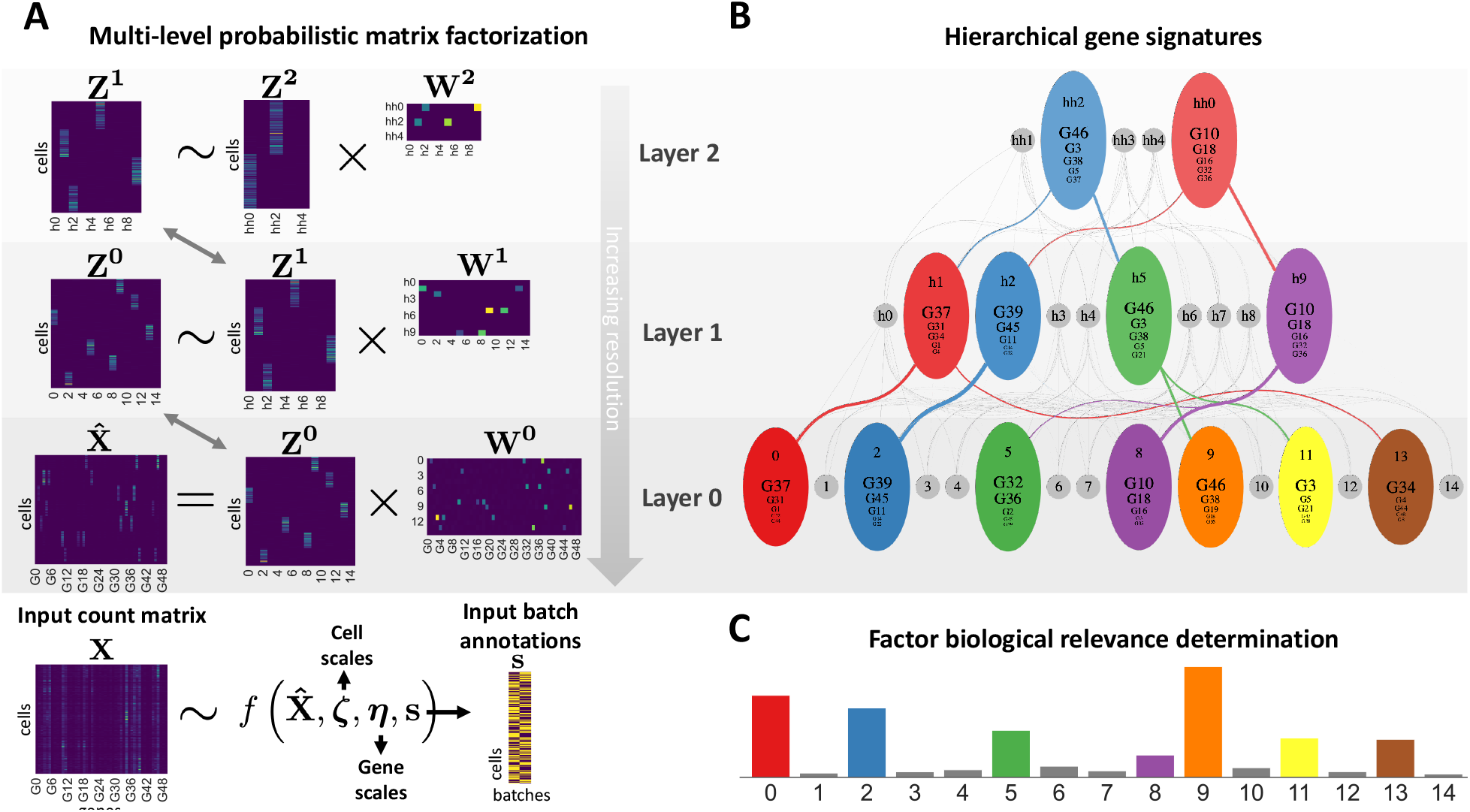
scDEF overview. (A) scDEF takes as input a matrix **X** of gene expression counts and learns a multi-level probabilistic non-negative matrix factorization model. The data **X** is modelled with a probability distribution *f* parameterized by a lower-rank approximation 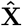, cell scales ***ζ***, gene scales ***η***, and optionally input batch annotations **s**. The approximation factorizes into non-negative matrices **Z**^0^ of cells-by-factors and **W**^0^ of factors-by-genes at the level of the highest resolution (layer 0). The matrix **Z**^0^ is also factorized, and this process repeats for the lower-resolution layers. (B) The model finds cell states in the data at multiple levels of resolution and identifies gene signatures for each one, as well as their hierarchical relationships. Each node in the graph corresponds to a factor in the hierarchy and shows the top 5 genes in the signature, with font sizes proportional to their scores. Gray nodes correspond to filtered out factors, with only the remaining ones for each layer colored. (C) scDEF automatically selects factors which capture biological instead of technical variation through the Biological Relevance Determination prior, creating a compact hierarchy.

At the highest level of resolution and at the lowest layer in the hierarchy, scDEF employs a novel prior distribution which favors factors that attribute a large weight to a small set of genes (Figure 1 C). This Biological Relevance Determination (BRD) prior leads scDEF to remove factors which affect many genes, which are unlikely to reflect biological signal but rather technical noise [11]. The model automatically selects the number of factors, favoring sparsity. Additionally, scDEF captures cell- and gene-specific scale factors in a similar fashion to scHPF [10], separating their influence from the multi-level factorization.

Optionally, scDEF may also take annotations indicating the batch of origin of each cell. This enables the model to capture batch-specific distributions of cell library sizes and gene detection rates. This is the only way in which batch effects are explicitly accounted for. The hierarchy and sparsity constraints further contribute to keeping only biological signal, resulting in robustness to technical batch effects while avoiding over-correction. The model may also be used in an informed manner, where gene sets are provided as input to guide the factors.

scDEF outputs a hierarchy of cell states (Figure 1 B) which can be used for dimensionality reduction, visualization, clustering, and gene signature identification. Due to its Bayesian nature, the model additionally provides posterior probabilities for all its parameters.

### Simulation study

We assessed the ability of scDEF to associate its factors with ground truth cell populations in single- and multi-batch simulations, using Splatter [24] to generate synthetic scRNA-seq data (Methods).

We compared scDEF with several alternative methods for cell population and gene signature identification (Methods) and evaluated them regarding their ability to recover the ground truth cell groups and gene signatures (Figure 2), as well as the separability of the cell groups in the latent space of each method and the quality of the hierarchies they learn (Supplementary Figure S1). The alternative methods comprise the standard approach based on PCA followed by Leiden clustering and Wilcoxon rank-sum testing, two other matrix factorization approaches (scHPF and NMF), and batch integration methods followed by Leiden and Wilcoxon rank-sum tests (Harmony, scVI, and Scanorama), which are the state-of-the-art methods for this task according to a recent benchmarking study [16]. In the multi-batch scenario, we consider scDEF both without batch information and with batch annotations (which is the default in the software).

**Figure 2:**
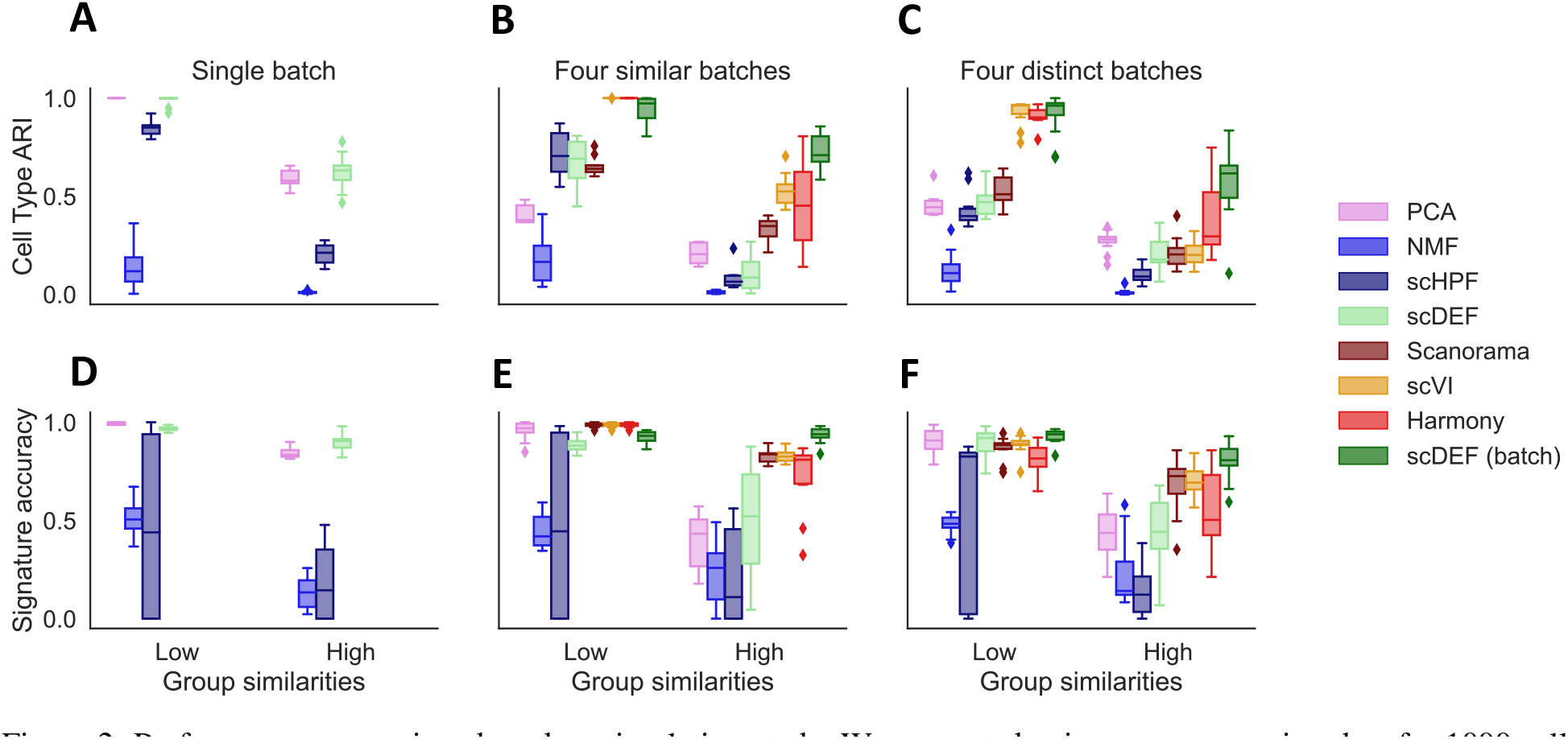
Performance comparison based on simulation study. We generated noisy gene expression data for 1000 cells per batch and 1750 genes containing 8 cell groups, and compared the ability of scDEF to various other methods in recovering the ground truth groups in single and multi-batch scenarios. We computed the adjusted rand index (ARI) between the simulated and inferred cell groups (Cell Type ARI, A-C), and the accuracy of the learned signatures (Signature accuracy, D-F), where higher is better. We varied the distance between the cell groups in all cases and simulated batches with shared cell groups (B, E) and also with batch-specific groups (C, F). For the single-batch setting (A,D), only the first four methods can be compared.

For each metric, we sort the simulation scenarios in Figure 2 from the easiest to the most difficult task, starting from the single batch case and ending with multiple batches including shared and batch-specific cell populations. For each case, we vary the similarity between cell groups, which we control through the scale of differential expression between groups (Methods).

We find that in the single-batch scenario, scDEF performs considerably better than the alternative matrix factorization methods and produces similar clustering and gene signature identification results as in the standard approach based on PCA and Leiden clustering (Figure 2 A). In terms of latent space, however, scDEF produces a low-dimensional representation which leads to a much stronger separation of the cell groups than any other approach (Figure S1 A), owing to its strong sparsity constraints. Additionally, by modelling a hierarchy directly, scDEF is able to recover the true underlying hierarchical structure better than the alternative methods (Supplementary Figure S1 D), which rely on multiple runs with different resolution parameters (Methods).

In the multi-batch scenario, we additionally benchmark scDEF against state-of-the-art integration methods (Figure 2 B, C, E, F and Supplementary Figure S1 B, C, E, F). Keeping the similarities between batches constant, we again vary the cell group similarities in both a scenario in which all batches are biologically similar, and another in which not all batches contain the same cell groups. In both cases, we find that scDEF without annotations leads to better results than both NMF and scHPF, highlighting the advantages of sparsity and a hierarchy in reducing technical batch effects. However, as expected, this is still outperformed by methods which aim to remove batch effects explicitly, especially if the cell groups are distinct enough to not be very confounded by the batch effects.

As the group similarities increase and batch effects become relatively stronger, providing annotations to scDEF leads to the overall best performance in comparison with all competing methods. This is even clearer for the case where the batches do not contain the same cell groups, where the performances of Scanorama, scVI, and Harmony become worse. Overall, in the multi-batch setting, with either poorly distinguished cell groups or with biologically different batches, scDEF is not only better at identifying the underlying cell groups, but also in reconstructing their signatures and hierarchical structure. Across all settings, scDEF detects cell states, their gene signatures, and their hierarchical structure more robustly than alternative methods.

### Analysis of scRNA-seq data from 3k PBMCs

We applied scDEF to the publicly available 3k PBMCs data set from 10x Genomics, which contains 2623 PBMCs from a healthy donor [25]. We ran scDEF with default parameters (Methods) and obtained a hierarchy containing three levels with 10, 5, and 4 factors for layers 0, 1, and 2, respectively, which enabled data visualization in 2D using UMAP embeddings and hierarchical signature discovery in an unsupervised manner (Supplementary Section S1). scDEF identified a compact hierarchy of cell states which is in agreement with the known PBMC cell subtypes, and whose gene signatures are sparse and include known marker genes for the populations they capture. In its informed versions (Supplementary Section S1.1), the model was able to assign cells to types based on input marker genes, find an upper level relationship between input subtypes, and resolve cell type-specific states.

This case study shows that scDEF can be used as a drop-in replacement to the usual scRNA-seq data analysis pipeline in a single-batch setting, enabling not only dimensionality reduction, unsupervised clustering, and gene signature identification, but also providing a meaningful hierarchy of cell states in the population we wish to study. Additionally, it provides, in the same model, the ability to use prior knowledge to guide the analysis, replacing the usual procedure in which users first assign cell types and perform clustering to identify higher-resolution states within coarse cell groups.

### Analysis of whole-animal scRNA-seq data

To further assess the ability of learning a hierarchy of cell states, we applied scDEF to scRNA-seq data from 21,612 cells from a whole adult animal, the flatworm *Schmidtea mediterranea*. This data set was generated and annotated by cell types in [26], which we use as ground truth to validate our results.

We first visualize the scDEF representations of cells at each layer in its hierarchy through UMAP embeddings where each cell is colored by the provided cell type annotations (Figure 3 A). Layer 0 strongly separates the differentiated cell types with neoblasts at the center, which is expected since they are the precursors of all cell types in the animal. In layer 1, cell types within the same tissue become less separated, with neurons, epidermal cells, muscle and parenchymal cells forming the main groups. At the upper layers, these groups are still visible but start to mix with the neoblasts in layer 2, and tissue types become even less clear in layer 3.

**Figure 3:**
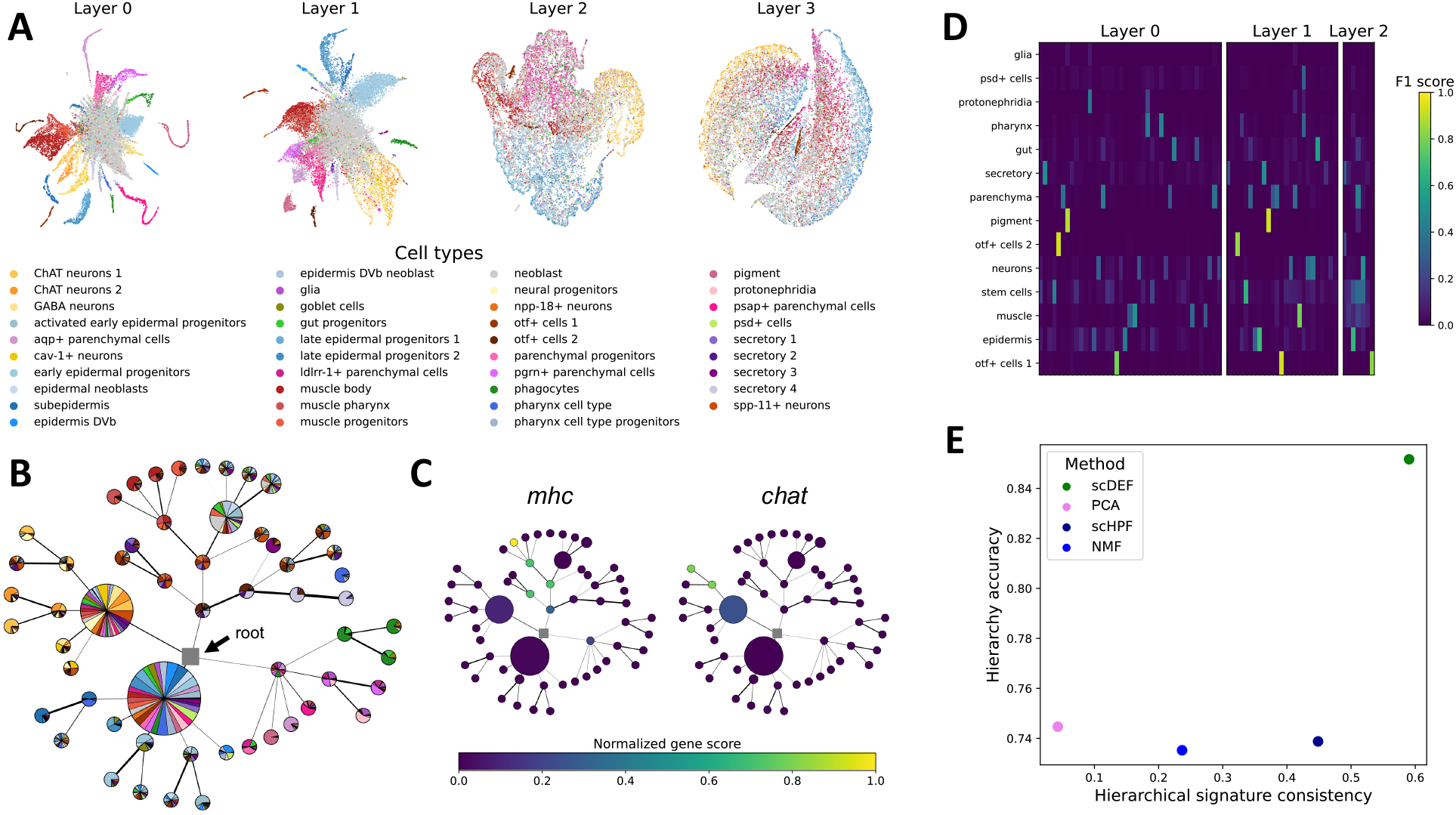
Application to a whole adult animal. (A) scDEF-based multi-layer UMAP embeddings of cells from the flatworm *Schmidtea mediterranea*. (B) scDEF hierarchy with nodes coloured by fraction of cells from each subtype in the organism, and scaled by proportion of cells attached to each factor. (C) scDEF hierarchy with nodes coloured by the normalized gene score of *mhc*, a known muscle-associated gene, and *chat*, a neuron subtype marker gene. (D) Association score between scDEF factors and coarse cell types. (E) Accuracy of the hierarchies learned by scDEF, Leiden, scHPF and NMF and the consistency of the signatures learned at each level.

The hierarchical structure is clear in the simplified tree (Figure 3 B), which we represent in a radial layout for compactness. We color nodes by the proportions of cell types that attach to each node, and scale the node sizes by the total number of cells that they contain. Clearly, nodes at the leaves correspond to more specific cell types and nodes near the root aggregate many cell types within the same tissue. For example, lineages corresponding to muscle cells and to neuronal cells are clearly visible in the top left region of the tree. scDEF directly learns gene scores for each factor at each level in the hierarchy, and we can map them onto the tree to identify genes that drive each lineage. In accordance with the original publication, we found that known planaria genes that are involved in muscle (gene *mhc*) and neuronal (gene *chat*) differentiation have a higher score in the lineages that are associated with cell types present in those tissues (Figure 3 C).

The association scores between factors at each layer and the coarse cell type annotations further confirms that scDEF learned a hierarchy in which cell types are captured at the lower layers and aggregated according to tissue type in the upper layers (Figure 3 D). Indeed, many factors in layer 0 associate with neurons, corresponding to different neuronal cell types, with fewer factors in layer 1 and finally only one in layer 2. The same observation applies to epidermal and parenchymal cells. For muscle cells, layer 1 already captures them strongly, highlighting the fact that muscle cells have fewer subtypes than, for example, neuronal cells. Cell types for which there is only one associated factor across the layers (for example, pigment cells) correspond to types for which the authors in the original publication did not identify a hierarchy, and indeed scDEF did not create a hierarchical structure for them either.

We compared the quality of the hierarchy learned by scDEF with the one obtained through a naïve method, in which we apply PCA (followed by Leiden), scHPF or NMF to obtain clusters at four levels of resolution and post-process them to identify the hierarchical structure learned across the different parameters (Methods). We then compare the cell group hierarchies they learned with the ground truth two-level hierarchy consisting of the main tissues at the higher level and cell types at the lower level. In addition, for each method we evaluate the consistency of the gene signatures at the different levels of the learned hierarchy. scDEF is the only method that jointly learns a hierarchy of cell groups and their gene signatures, and it performs better than the alternatives in both metrics (Figure 3 E). By modelling the presence of a hierarchy directly, scDEF bypasses the need to re-cluster certain regions where a higher level of detail is required, and instead directly provides different levels of resolution, their relationships, and their gene signatures.

### scDEF performs robust batch integration

As the number of scRNA-seq data sets increases and both their experimental and biological conditions vary, methods that are able to jointly analyse multiple data sets and remove technical differences while keeping the biological effects that are specific to each batch become increasingly relevant. In contrast to the current paradigm in which batch integration methods aim to maximize the similarity between batches, scDEF instead aims to produce factors that highlight biological signal and remove technical variation. This is achieved via a combination of sparse factors in a hierarchical structure, and capturing batch-specific library sizes and gene detection rates. Here, we apply scDEF to a setting with biologically similar batches and another with batch-specific biological signal and show that scDEF performs well in both situations. We show the main results from both data sets side by side in Figure 4 to make the differences clear.

**Figure 4:**
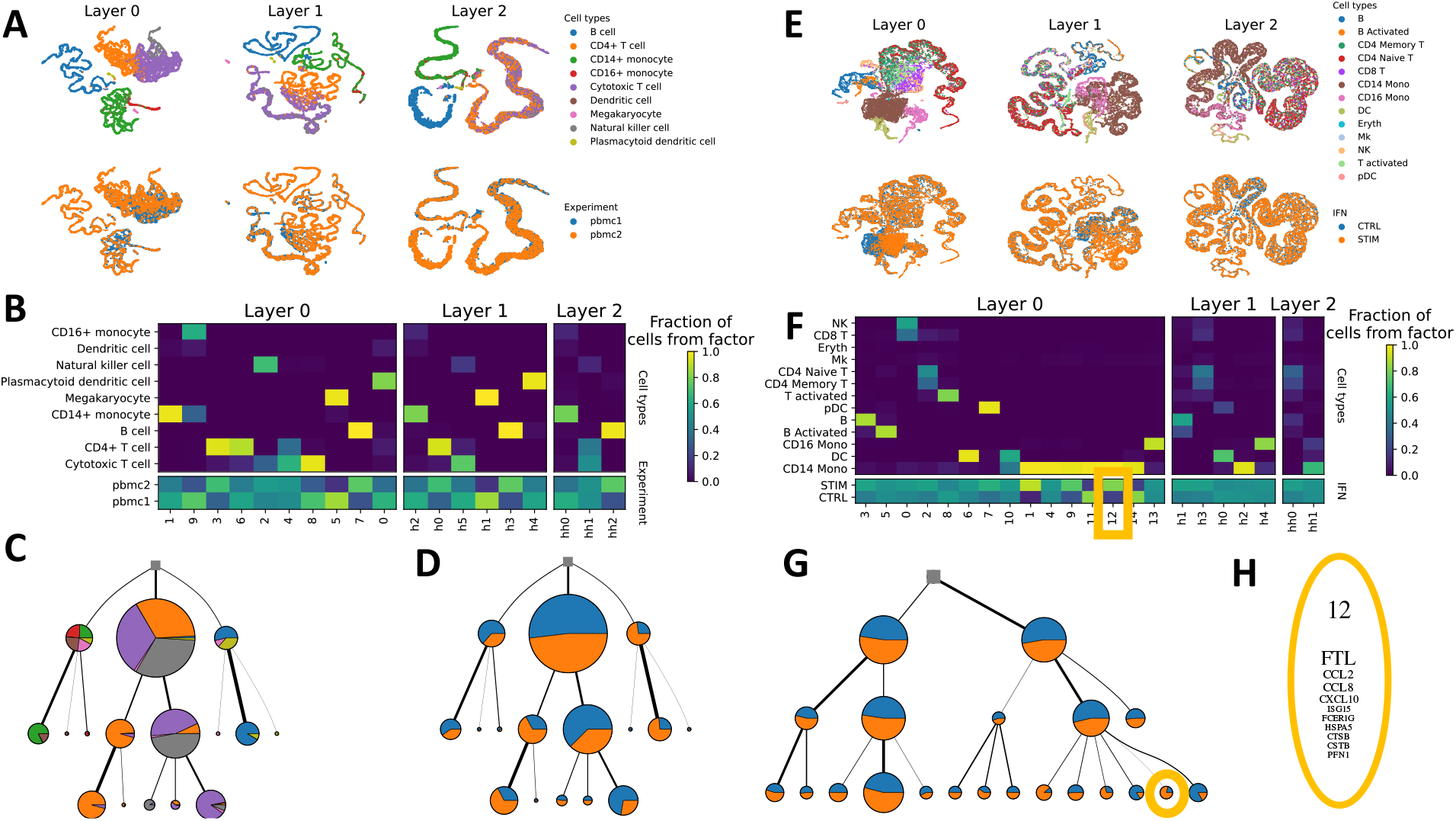
Application to data containing healthy PBMCs from two batches (A-D) and one batch of interferon-stimulated PBMCs and another containing unstimulated PBMCs (E-H). (A,E) scDEF-based multi-layer UMAP embeddings of cells colored by their given cell types and batch of origin. (B,F) Fraction of cells in each factor from scDEF that belong to each cell type and batch annotation. (C, D, G) Learned cell state hierarchies coloured by fraction of attached cells from each cell type and batch. (H) Highlighted factor 12 contains mostly interferon-stimulated cells and a gene signature for response to interferon stimulation.

First, we applied scDEF to a subset of data from the Broad Institute PBMC Systematic Comparative Analysis study [27], consisting of two batches of healthy PBMCs sequenced separately with 10x Chromium v2 (“10x Chromium v2 A” and “10x Chromium v2 B”) and containing 3222 and 3362 cells. We expect them to mostly consist of the same cell types, and so nearly all batch-specific variation to be due to technical effects that we wish to remove. Indeed, jointly performing a UMAP projection of all cells from both data sets without any correction reveals very strong batch effects (Supplementary Figure S2 A, B).

scDEF learned a hierarchy with three layers (Figure 4 A-D, Supplementary Figure S3). To visualize the structure of the data at each layer, we map the weights of each cell to each of that layer’s factors to two dimensions using UMAP, and color the cells by both cell type annotations and batch (Figure 4 A). We observe a progressive mixing of cell types from the lower to the upper layers, while maintaining a consistent mix of cells from all batches across all the layers, indicating successful batch integration.

The association between the cells attached to each factor at each layer and the cell type annotations further confirms this, meaning that the factors can be used as batch-corrected clusters (Figure 4 B). Indeed, there are clear associations between factors and cells types, with upper layers capturing meaningful groups (e.g., factor hh0 of layer 1 captures all monocytes, and factor hh1 captures all T cells), but no stronger association of any particular factor over another in any layer with batch. The fact that the hierarchy captures cell type information instead of batch-specific biases is also clear from the tree representation (Figure 4 C, D), in which cell types are aggregated according to known functional relationships, and there is no node with a strong over-representation of either batch.

By default, scDEF uses batch annotations to correct for batch-specific gene detection rates, and this was the case for the results shown above. Correcting for this per-batch, per-gene effects is the only way in which batch annotations are used in the model, which is able to further correct for technical variation due to its sparsity and hierarchy constraints. The power of these two crucial aspects of scDEF is clear from the results on the same data where no batch annotations were provided to the model (Supplementary Figure S2 D). Most populations are still identified irrespective of batch, and only cytotoxic T cells still show batch-specific effects. In contrast, the usual approach without integration based on Leiden identifies clusters which are all batch-specific (Supplementary Figure S2 C). Remarkably, the upper levels of the scDEF hierarchy capture the same coarse cell groups regardless of whether batch annotations were provided (Supplementary Figure S2 D, E). We compared both scDEF versions (without and with batch annotations) with competing methods to identify cell populations, including standard batch integration tools like Harmony, scVI and scanorama (Supplementary Figure S2 F). scDEF without batch annotations is not able to perform batch integration as accurately as those methods, but it is still much better than any other method that does not use those annotations. However, in the default mode with batch annotations, scDEF is competitive with the state of the art in this task.

We then applied scDEF to a dataset consisting of PBMCs from two lupus patients, one of whom was treated with interferon, to examine cell type-specific responses to the stimulation. The data were generated in [28] using a droplet-based protocol. When the batches come from different conditions, common batch integration methods will fail to resolve their differences, which may be biologically relevant. We expect cell type-specific reactions to the stimulus, which means that scDEF should be able to identify factors indicating the genes that become active upon this stimulus. In particular, the authors reported a monocyte-specific increase in the expression of *CXCL10* and *CCL8*.

As in the previous example, a batch-uncorrected UMAP embedding show a clear separation between batches, and these strongly drive the basic Leiden clustering results (Supplementary Figure S4 A B, C). In contrast, the UMAPs based on the four layers that scDEF learned show a clear separation between cell types and only a monocyte-specific split between the different conditions (Figure 4 E). In particular, while factors 1, 4, 9, 11, 12, and 14 at layer 0 all contain only CD14 monocytes, some of them have either these cells from the CTRL (11, 14) or the STIM (1, 9, 12) batch. In other words, factors 1, 9, and at 12 at layer 0 strongly associate with monocytes from the stimulated batch, whereas the remaining factors are well mixed between batches and associate with the other cell types (Figure 4 F). The hierarchy additionally shows that scDEF learned meaningful cell type associations and found canonical marker genes for subpopulations (Supplementary Figure S5). If we color the hierarchy by the proportion of cells from each batch, we identify only a few nodes in the lowest layer in which either batch is over-represented, in accordance to the heatmap (Figure 4 G). The signature of the stimulation-specific factor 12 contains genes previously described to be related to monocyte-specific response to interferon, including *CCL8* and *CXCL10*, and also *ISG15* (interferon-stimulated gene 15) (Figure 4 H). We thus confirm that scDEF was able to simultaneously identify all the cell types, the monocyte-specific response to interferon, and the corresponding gene signatures.

We also assessed the case in which no batch annotations were given to scDEF and found that it captured more batch-specific factors than when annotations were provided (Supplementary Figure S4 D, E). In particular, while most cell types were well integrated between batches, more factors separated the stimulated monocytes. This difference propagated to layer 1, which instead of containing a hierarchical factor for monocytes as before, contained one per monocytes in the different conditions. If we set as ground truth annotations the cell types with monocytes separating between stimulated and controls, we observe that the performance of scDEF, regardless of using batch annotations, is better than any other batch integration and clustering method combination, which always remove the true batch-specific signal in the data (Supplementary Figure S4 F). These results illustrate that the hierarchical structure and the sparsity constraints in scDEF enable it to fix most technical batch effects while keeping biological variation intact. Nevertheless, using batch annotations yields stronger batch correction while still keeping the biological variation, even though its effect may be slightly reduced. scDEF thus provides a balance between no batch correction, where all populations are batch-specific, and over-correction, where there are no batch-specific populations.

### scDEF identifies shared and patient-specific variation in a high-grade serous ovarian cancer cohort

Cancer is a disease hallmarked by heterogeneity both across and within patients. The design of effective therapies may benefit from a characterization of which cancer cell populations are specific to each patient, and which are shared across them. To perform this distinction, typical batch integration methods suffer from over-correction problems, where patient-specific populations are lost in the integration [22]. However, without any correction, technical batch effects may hide shared gene expression profiles. We tested whether scDEF could recover shared and patient-specific populations in a HGSOC data set containing 11 patients with samples from before and after chemotherapy [29].

We applied scDEF to this data set providing the model with patient IDs as annotations. scDEF learned a hierarchy with 3 layers containing 27, 7, and 2 factors (Figure 5 A, B). Both factors at the top layer contain a mixture of cells from all patients, indicating that there is a hierarchy of cell states which is shared across the cohort. However, lower levels in the tree reveal factors which are patient-specific, in particular corresponding to cell populations that are present within the patients with the largest number of cells, EOC136 (with 2178 cells) and EOC733 (with 1754 cells). Interestingly, patient EOC733 has two subpopulations, one in either side of the main split of the hierarchy (factors 5 and 24), indicating that this patient’s tumor may be particularly heterogeneous.

**Figure 5:**
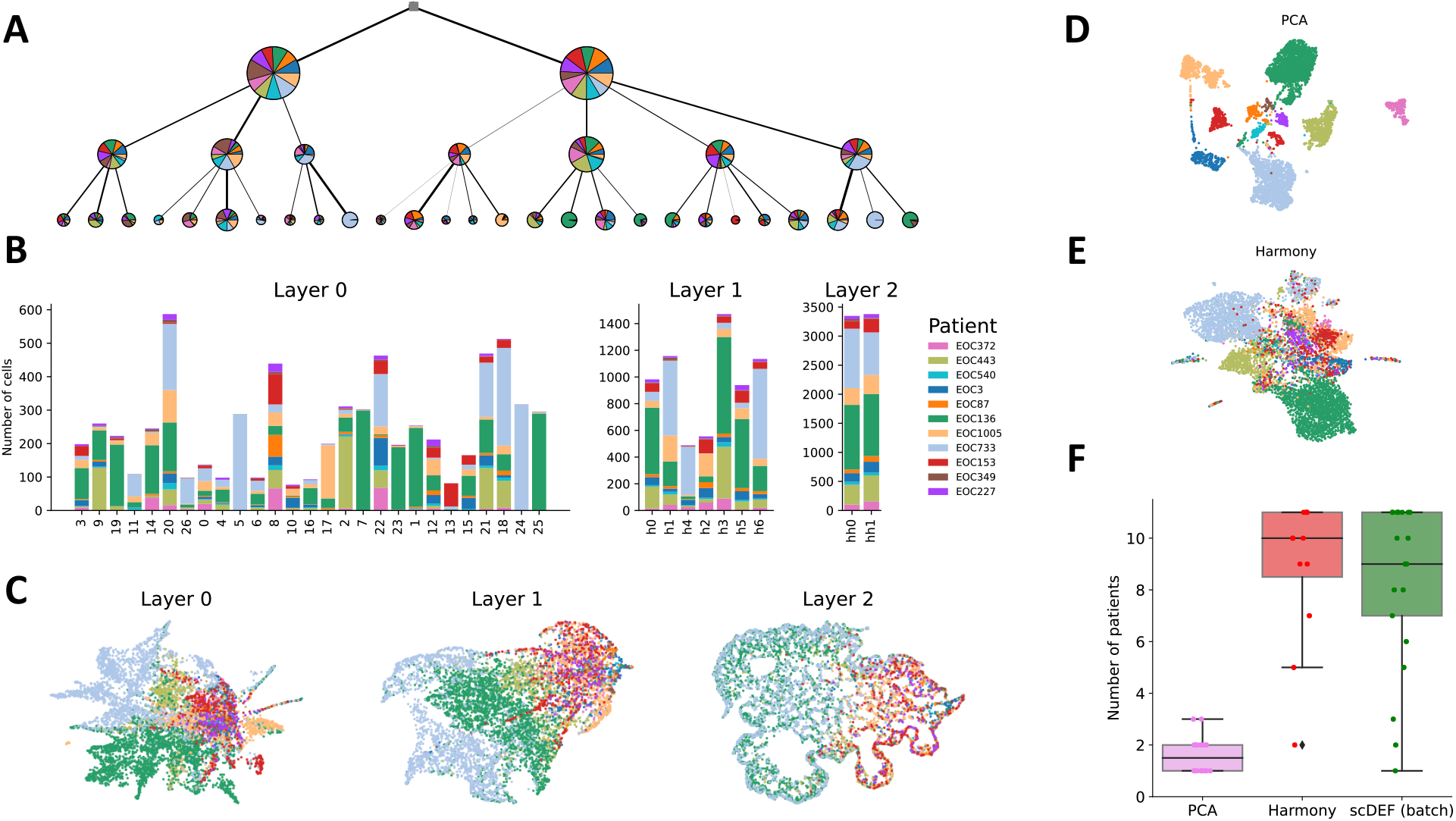
Application to a cohort of high-grade serous ovarian cancer patients. (A) Hierarchy learned by scDEF with nodes coloured according to proportion of cells from each patient and area scaled by the number of cells. (B) Number of cells from each patient in each factor in scDEF. (C) UMAP embeddings of the scDEF layers with cells colored by patient. (D) UMAP embedding based on PCA dimensionality reduction. (E) UMAP embedding based on Harmony batch correction. (F) Number of patients in each cluster obtained from Leiden on PCA and Harmony, and in each factor obtained by scDEF.

The UMAP representation of the cell weights in layer 0 shows a mixture of many cells from different patients except patients EOC136 and EOC733, which mostly separate from the rest (Figure 5 C). However, even some groups of cells from these patients mix in with cells from other patients, which is also visible from the proportions of cells from each patient in factor (Figure 5 B). For example, about half the cells in factor 9 are from EOC136, and the other half from EOC443. The majority of cells in factor 2 correspond to patient EOC443, and about 20% come from patient EOC136. The upper layers keep patients EOC136 and EOC733 separate while showing sets of cells from other patients grouping together, with the top layer showing one main split between the two main patients and the remaining ones. However, as indicated from the cell to factor assignments (Figure 5 B), this split does not correspond to the main split into two factors at layer 2. Instead, this indicates that within each of these two factors, patients EOC136 and EOC733 have different weight distributions than the other patients. The factor-based split in this UMAP is therefore vertical and not horizontal (Figure 5 C).

In contrast, an uncorrected UMAP produces a strong separation of cells from all patients (Figure 5 D), while a Harmony-corrected representation shows an approximation of cells from different patients (Figure 5 E). scDEF (with batch annotations provided as input) provides an intermediate solution to these two extreme approaches that is closer to Harmony, but keeps more effects which are specific to patients in addition to effects that are shared across them. By running the Leiden clustering algorithm on both the unintegrated space and the Harmony-corrected representation, we may compare these three methods in terms of their ability to find both cell populations which are patient-specific and populations which contain cells from many patients (Figure 5 F). As expected, the approach without integration (PCA) tends to produce only patient-specific clusters, due to the strong presence of batch effects. At the opposite end of the spectrum lies Harmony, for which the majority of clusters include almost all 11 patients. scDEF, on the other hand, produces a majority of clusters which are shared across many patients, but has a long tail of clusters with subsets of patients. scDEF can be interpreted as a compromise between no correction and strong batch correction, erring on the side of correction when batch annotations are provided (which is the default setting).

We then sought to find whether the hierarchical structure captured differences between treatment-naive and post–neoadjuvant chemotherapy (post-NACT) cells. We found that scDEF not only identified factors that contained cells either only from treatment naive or treated samples, but also factors that contained mixtures of these two groups, indicating shared variation or transient states (Figure 6). In addition, scDEF found factors that contained either mostly untreated or mostly treated cells, even though they belonged to multiple different patients. For example, in layer 0, factors 12, 10 and 15 are enriched for cells from mostly post-treatment samples from multiple patients. In layer 1, this is the case for h2, h0 and h5.

**Figure 6:**
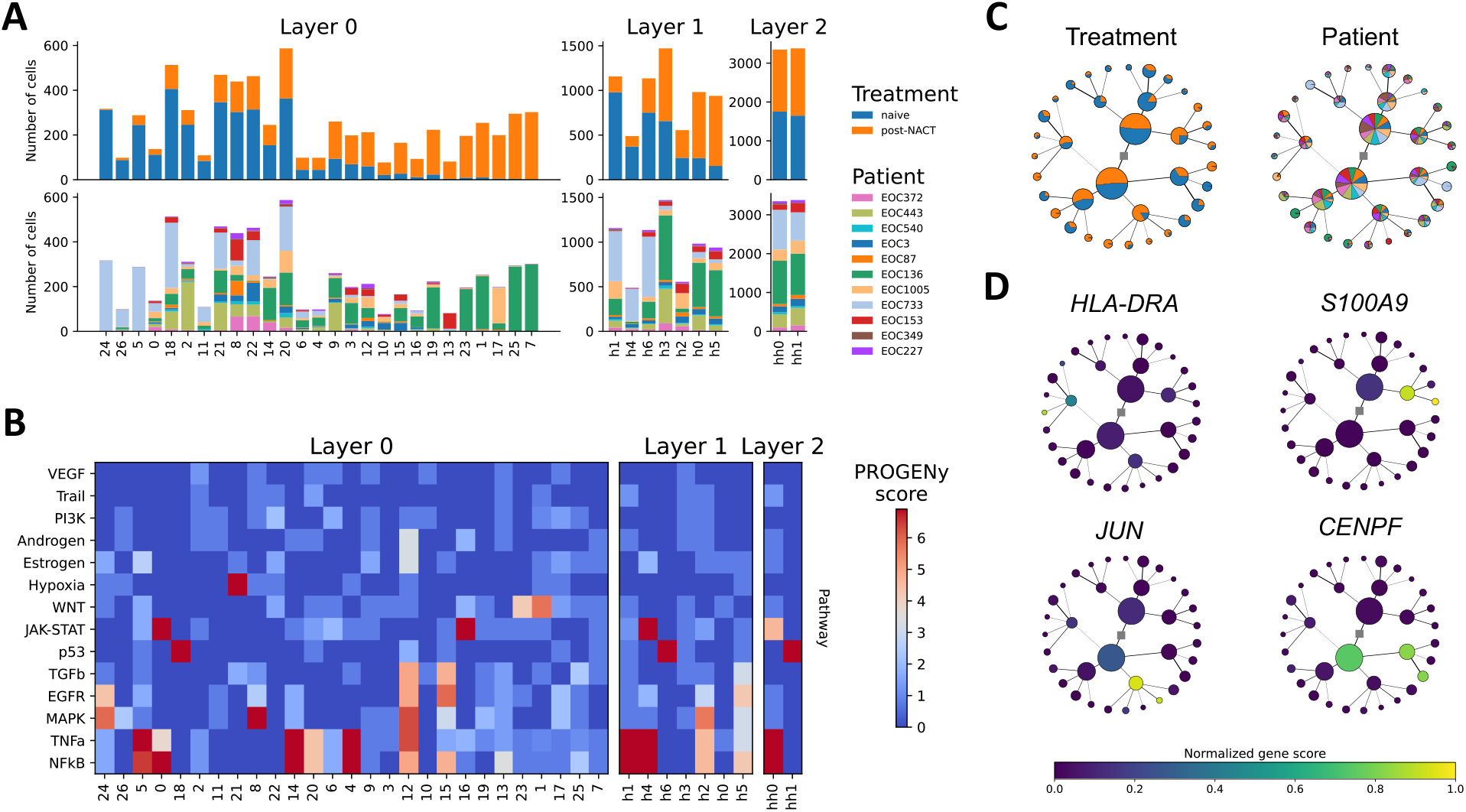
Application to a cohort of high-grade serous ovarian cancer patients continued. (A) Number of cells from each patient in each factor in scDEF, coloured by treatment status and patient ID, and sorted from lower to higher fraction of post-chemotherapy cells. (B) PROGENy pathway activities per scDEF factor, with factors sorted as in panel A. (C) Radial representation of the hierarchy with nodes coloured by treatment status and patient ID, and by (D) normalized score of selected genes in the gene signatures in the factors of the hierarchy.

In order to interpret the functional meaning of the hierarchy, we used decoupler [30] and PROGENy [31] to perform over-representation analysis on the gene signatures in each factor of the learned scDEF hierarchy (Figure 6 B, Methods). At the top level (layer 2), one factor is only enriched for the p53 pathway, and the other shows strong association with immune-related pathways (NFKb, TNFa, JAK-STAT). At the next level of resolution (layer 1), we find that factor h5, which is enriched for post-treatment samples, has a large score for the EGFR pathway, and h2 shows a high activity of the MAPK pathway. In layer 0, factors 12 and 15 exhibit a high association with TGFb and EGFR, which are associated with proliferative phenotypes, but also a high score for NFKb, which is associated with immune response and cell survival.

We additionally highlight a few of the top scoring genes for different parts of the hierarchy (the top 10 genes for each factor in the hierarchy are shown in Supplementary Figure S6). We found that the factor with the highest number of cells (factor 20) is strongly enriched for genes related to MHC class II antigen presentation (e.g. gene *HLA-DRA*, Figure 6 D), and that this signature includes pre and post-treatment samples in nearly the same proportion, indicating a conserved sub-population regardless of patient and treatment. Additionally, we find genes associated with cell cycle progression and differentiation (e.g. *S100A9*) in factor h0.

Finally, we observed that factors h5 and h6 both contained genes associated with stress response. For example, factor h5 attributes a high score to *JUN* and *FOS*, which tend to be over-expressed in cells under stress, and factor h6 to *CENPF* and *CENPA*, whose expression is associated with cell cycle and the p53 pathway. Strikingly, despite being closely related in the scDEF hierarchy, factor h5 contains mostly cells from post-treatment samples and h6 from pre-treatment (Supplementary Figure S6, Figure 6 A). This finding suggests a transient state induced by chemotherapy, in which cells with strong activity of the p53 pathway are better equipped to survive through treatment.

## Discussion

Identifying cell populations and their gene signatures are key tasks in single-cell data analysis. Conventional approaches rely on a two-step procedure in which cells are clustered and differential expression tests are carried out between the clusters. These approaches ignore the fact that cell states are often arranged in a hierarchy, and require extra pre-processing steps to remove technical batch effects. Here we have followed a different approach based on a multi-level Bayesian matrix factorization model which addresses both tasks jointly and does not suffer from those limitations. The multi-level factorization ensures not only that we are able to learn hierarchical and sparse gene signatures, but also that those gene signatures are more robust to noise than a single-level counterpart. This is useful in single-batch scenarios, but its benefits are even clearer in multi-batch applications. With the constrained factorization, technical batch correction can be achieved using only a global correction for the differing gene detection rates for each batch.

Our results show that scDEF is able to obtain representations of the data which respect meaningful biological hierarchies in both single and multi-batch scenarios. In particular, scDEF recapitulated known cell type hierarchies in PBMCs and a whole adult animal. In a cohort of cancer cells from multiple patients, where cell types are not clearly defined, scDEF found hierarchies of cell state that describe differences between treatment naïve and post chemotherapy samples that are consistent with previous findings. This is a particularly challenging task where batch is confounded with biological conditions [32], but scDEF was able to identify both shared and patient-specific variation, indicating that it did not over-correct the signal.

We have defined scDEF using a sparse gamma deep exponential family with cell- and gene-specific size factors and the novel BRD prior for automatic model selection. While the architecture of this model is akin to a multi-layer neural network, by treating it as a generative model and performing approximate Bayesian inference we are able to include interpretable constraints such as non-negativity and sparsity in a probabilistic framework. This approach has advantages over models such as LDVAE [33], which is a variational autoencoder-based model without non-negativity and sparsity constraints, which leads to denser and less interpretable gene signatures, as well as to not promoting cell clustering. However, because we approximate the posterior distribution of all variables in our model, we are unable to make use of amortized inference techniques such as the ones employed by LDVAE and scVI. This makes scDEF generally slower than these methods.

Compared to supervised approaches, which are increasingly used to identify cell populations in single-cell data, scDEF does not require either cell annotations, nor input gene sets, which may limit our discoveries if too limited. This is of particular importance in cancer settings where the evolutionary process that leads to tumor growth is largely unpredictable and generates highly heterogeneous cell populations. By finding unknown cell states and their gene programs robustly across data sets, scDEF is an appropriate tool for the analysis of large-scale cancer studies, which are increasingly common and hold the potential of contributing to the design of personalized treatments [34].

In general, matrix-factorization methods have been gaining increasing attention in single-cell data analysis due to their interpretability. scDEF is the first of these methods to simultaneously model a hierarchy of factors and to introduce the BRD prior to determine the number of biologically meaningful factors. By doing this in a Bayesian framework, scDEF directly provides confidence estimates on the gene signatures it infers from data. In addition, the model can be used in both a fully unsupervised manner or with prior gene sets, making it suitable for a wide range of scenarios. In the age of widespread single-cell data studies to discover, track, compare, and characterize cell populations, scDEF is well positioned to be an important tool in robustly learning gene signatures across data sets.

## Methods

### scDEF model

scDEF models gene expression counts as a multi-layer Bayesian sparse non-negative matrix factorization. Let **X** be the input gene expression counts matrix of size *C* cells by *G* genes, and let *L* be the number of layers, where each layer *l* has *K*_*l*_ factors. If the data come from different batches, let *B* be the number of batches, and let *s*_*cb*_ = 1 if cell *c* belongs to batch *b*, and 0 otherwise. scDEF is defined through a generative process which describes how data arise from the model, outlined in Algorithm 1.

We describe the model starting from the likelihood and going backwards through the generative process. The observations 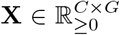 are obtained from the matrix product **Z**^0^**W**^0^, which is a rank-*K*_0_ approximation to **X**, with 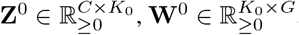, and *K*_0_ < *G*. Specifically, the observed gene expression *x*_*cg*_ of gene *g* in cell *c* is drawn from a Poisson distribution with mean defined by the vector product between row *c* of **Z**^0^ and column *g* of **W**^0^. Each cell *c* has weight 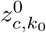 for factor *k*_0_, and factor *k*_0_ gives weight 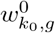 to gene *g*. To ensure non-negativity, 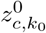 and 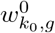 are drawn from Gamma distributions. This is the Poisson matrix factorization model [35], and it expresses the idea that the gene expression profile of each cell is the product of multiple gene programs, each containing a small set of genes.

In addition, **Z**^0^ is itself parameterized by a rank-*K*_1_ approximation **Z**^1^**W**^1^, with 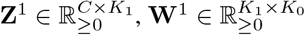, and *K*_1_ < *K*_0_. Whereas **W**^0^ groups the *G* genes into *K*_0_ factors, **W**^1^ groups the *K*_0_ factors into *K*_1_ higher-level factors. Similarly, **Z**^1^ indicates which of the *K*_1_ factors best describe each cell *c*. Figure 1 B illustrates this: the lower *K*_0_ circles represent the highest resolution factors, with the top-5 genes with highest weight **W**^0^ indicated inside each one. The upper *K*_1_ circles represent the higher-level factors, and the thickness of the edges connecting both layers represents the weights **W**^1^ that the upper-level factors attribute to each lower-level factor. This structure is repeated for *L* layers.

#### Algorithm 1

scDEF generative process

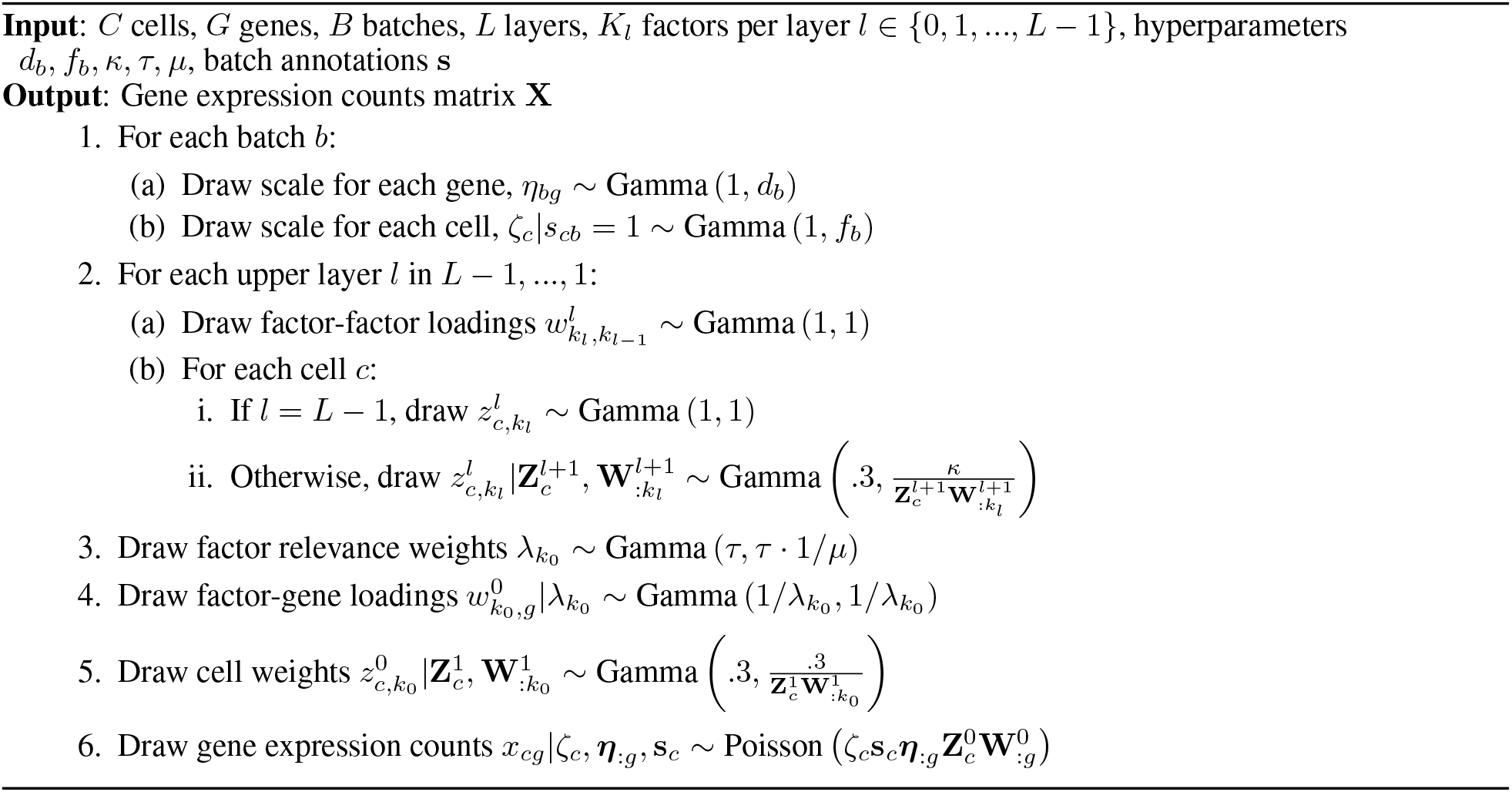

We use the observation [11] that biologically relevant factors affect few genes, leading to sparse profiles, in contrast to technical factors which affect many genes. The factor weights at layer 0 are modelled by the Biological Relevance Determination (BRD) prior, in analogy to the commonly used Automatic Relevance Determination (ARD) prior. The ARD is used to automatically drive the weights of factors that are not used to 0, retaining only those factors that explain a sufficiently large portion of the data. Our BRD prior follows this idea but instead will drive non-sparse factors to irrelevance, thus keeping only the biologically relevant ones. The factor relevance weights are drawn from a Gamma distribution centered at *µ* with a strong concentration *τ* and interact with the factor weights by modelling their concentrations. If *λ* is large, the factor is sparse owing to its peak at 0 and long tail. In contrast, if *λ* is small, then the gene weights of that factor tend to be large and not sparse. This enables us to ignore factors with small importance. In practice, this means we can run scDEF with large *K*_*l*_, and the BRD prior will automatically turn off non-sparse factors at layer 0, and that selection will propagate to the factors in the higher layers. By default, scDEF uses 4 layers with 100, 60, 30, and 10 factors, and we set *τ* = 10^3^ and *µ* = 10^2^.

To account for cell- and gene-specific scale factors, which are quantities that may confound our analysis of gene signatures and cell states, we explicitly model them in ***ζ*** and ***η***, respectively. Following [10], the hyperparameters *d* and *f* of the distributions of these random variables are set to preserve the empirical variance-to-mean ratio of the total molecules per cell or gene, in each batch:

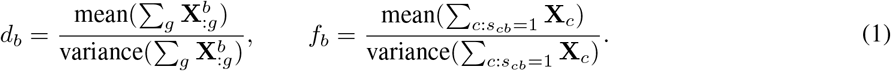

Modelling these quantities as being specific to each batch is the only direct way in which batch information is given to the model. scDEF can be seen as an extension of scHPF with higher layers, the BRD prior, and batch-aware scale factors for both cells and genes.

Finally, we set the hyperparameter *κ* according to the average number of detected genes across cells. If this number is high, then each cell strongly informs the model, and *κ* may be low. Otherwise, we require a larger *κ* to ensure the hierarchy is used. As a simple rule, we set *κ* to 1 if the average number of detected genes is over 10^3^, and to 10 otherwise.

### Using prior information

We now modify the scDEF model above to enable the use of known gene sets to guide the factorization, depending on the resolution of those sets. We name this version informed scDEF (iscDEF). It can be used in two distinct modes.

Mode 1 simply sets the lower layer factors to prefer genes from specific gene sets, with one factor per gene set, and learns a higher-level structure. The model is overall the same as scDEF, with changes in the prior for the gene signatures **W**^0^. We also remove the BRD prior, as the input high-resolution gene sets are assumed to be present in the data. The prior for **W**^0^ is therefore set as

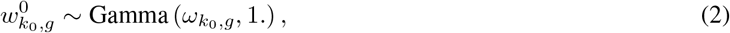

where 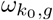 is the prior weight that gene *g* has in factor *k*_0_, which we collect in a matrix **Ω**. If *g* belongs to the gene set that we wish to encode in factor *k*_0_, then 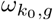 is set to 10. Otherwise, it is set to 1.

Mode 2 associates the gene sets with the higher layer and learns gene set-specific substructure in the lower layers. Assuming we have *P* gene sets and *L* layers, we create an iscDEF where layer *L* − 1 has size *P* and each subsequent layer *l* has size *PM* (*L* − *l*), where *M* is a hyperparameter determining the number of new factors per layer per gene set. We then set the prior for each **W**^*l*^, *l* > 0 as

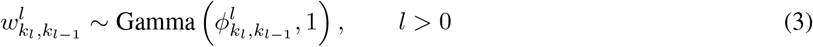

where 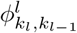 is the prior weight that factor *k*_*l*_ gives to factor *k*_*l*−1_ in the next layer down the iscDEF. This will promote each higher level gene set to find its own specific substructure. The gene set information is again used in the prior for **W**^0^, but in this case each one of the *ML* factors per gene set in this layer have the same prior, according to the gene set at the top. This structure is again encoded in the *K*_0_ *× G* matrix **Ω**. In this mode we keep the BRD prior to remove unnecessary lower-level factors. The prior for **W**^0^ is then

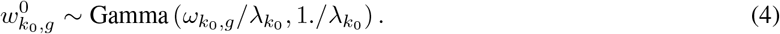

### scDEF inference

Given UMI count data **X**, we wish to find the scDEF model parameters defined in Algorithm 1, that best describe the data. We pursue a Bayesian approach and target the posterior distribution over all variables in the model, *P* (**Z, W, *λ, ζ, η***|**X**). Since the model is non-conjugate, this distribution is not analytically tractable. In this case, the two main approaches to find the posterior distribution are sampling methods, such as Markov chain Monte Carlo (MCMC) [36], and variational inference [37]. Here we use variational methods, because they are often able to obtain an approximation faster than MCMC and are generally better suited to large data sets [38]

In variational inference, we posit a simpler family of distributions over the latent variables of the model, *Q*(**Z, W, *λ, ζ, η***), the parameters of which are determined by minimizing the Kullback-Leibler divergence between *Q*(**Z, W, *λ, ζ, η***) and the true posterior *P* (**Z, W, *λ, ζ, η***|**X**) or, equivalently, by maximizing the evidence lower bound (ELBO),

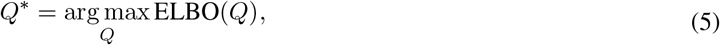

with

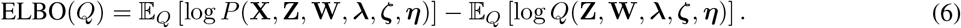

We use a mean-field approximation for *Q* in which the posterior distribution over the latent variables factorizes completely, and we let each marginal variational distribution be in the same family as the prior of the corresponding latent variable (which in our model means that they are all approximated with Gamma distributions), with its own variational parameters:

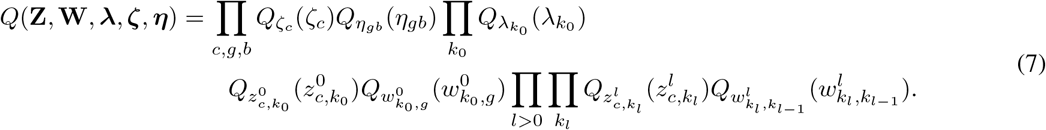

This approach is also used in [10]. However, since our model is not even conditionally conjugate due to the upper layer, we can not compute the ELBO analytically and may not use analytical updates for the variational parameters. We thus resort to black-box variational inference [39, 40], in which the variational parameters are updated using Monte Carlo approximations of their gradients with respect to the ELBO. Collapsing all the latent variables into ***θ*** for readability, we then approximate the ELBO as

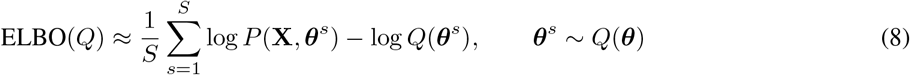

where *S* denotes the number of Monte Carlo samples from the variational distribution. We employ automatic differentiation with JAX [41] to compute the gradient of all the variational parameters with respect to the ELBO, and use the widely adopted gradient descent method Adam [42]. In practice, we run scDEF with *S* = 10, which enables a good trade-off between speed and low gradient variance. By default we do not use minibatch optimization, but this is supported by our software. There, we sample, at each iteration, a subset of cells and optimize only their local variational parameters, and scale the gradient with respect to the global variational parameters accordingly. We initialize the Gamma variational distributions by setting their shapes and rates to their prior values.

### Filtering factors

The BRD prior automatically leads the learned scDEF towards using only a small fraction of the total possible number of factors at layer 0. At each layer, we attach each cell to the factor that it assigns the most weight to, and we remove factors that have fewer than 10 cells attached. Additionally, we only keep factors in layer 0 which have a posterior BRD (*λ*) mean that is at least 3 times the inter-quartile range of all the posterior BRD means.

### Extracting a tree from the learned hierarchy

The learned scDEF corresponds to a hierarchical network where each pair of adjacent layers is fully connected. However, the prior distributions on the connection weights **W** lead to sparsity, which we may leverage to remove weak connections and simplify the representation of the hierarchy (Figure 1 B). In doing so, some factors may end up having only one parent, and we further simplify the hierarchy by collapsing them into one. This leads to a tree representation of the learned scDEF hierarchy.

### Cell embeddings per layer

Each layer in the learned scDEF can be used as a low-dimensional embedding that we can use for the visualization of cells in two dimensions using UMAP projections as is commonly done in scRNA-seq data analysis. We use the scanpy Python package to compute a k-nearest neighbours graph based on the scDEF factors with sc.pp.neighbours and the sc.tl.umap function to visualize the embeddings in two dimensions, with default parameters. Because in scDEF these cell embeddings **Z** have very skewed distributions owing to their sparse gamma priors, and a lot of cells attribute strong weights to one factor and very little to the rest, their 2D projections often form a line, which is a natural consequence of the model.

### Factor association score with cell annotations

We compare the cell-to-factor assignments with user-provided cell annotations by computing the F1 score. For plotting, we sort the cell annotations in the rows of the matrix containing these scores across all scDEF layers using Ward clustering.

### Gene signature identification

We define a gene signature as a truncated ranked list of genes in each factor, where by default we use the top 20 genes with the highest score. For layer 0, the rankings are obtained directly from the posterior mean of **W**^0^. For layer 1, they are obtained by the inner product of the posterior means of **W**^1^ and **W**^0^. In general, for layer *l*, the gene scores for each factor *k*_*l*_, which we denote 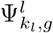 are obtained recursively as

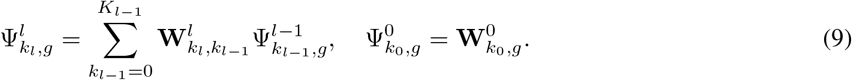

To assess the stability of the signatures we take 100 Monte Carlo samples from the variational distribution of **W** and, for each sample, create the gene signatures for all factors in all layers as described previously. The Jaccard index for each signature across all posterior samples then indicates the posterior stability of each signature.

### Pathway enrichment analysis

For each gene signature, we perform an over-representation analysis using the decoupler method [30] and the PROGENy database [31], which contains the top 500 genes associated with 12 pathways: Androgen, EGFR, Estrogen, Hypoxia, JAK-STAT, MAPK, NFkB, p53, PI3K, TGFb, TNFa, Trail, VEGF and WNT. For each factor, we report the z-scores computed across all pathways.

### Alternative methods

We compared scDEF to the standard clustering and marker gene identification method based on Leiden clustering followed by differential expression analysis, two other matrix factorization methods, NMF and scHPF, and three of the top-performing batch integration methods according to a recent benchmarking study [16]: scVI, Harmony, and scanorama. We also applied Leiden and computed differential expression to the results of the integration methods, since these simply produce batch-corrected embeddings.

- **PCA**: We ran the usual scRNA-seq data processing steps based on highly variable gene filtering, log-normalization, library size and mitochondrial content effects removal, scaling, followed by dimensionality reduction with PCA, neighborhood graph construction and Leiden clustering. For clustering with Leiden, we used resolution parameters of 1.0, 0.6, 0.3, and 0.1, following the layer sizes used by scDEF. We used the implementation provided in the scanpy Python package.
- **NMF**: We ran non-negative matrix factorization on the normalized counts data using 30, 20, 10, and 5 factors to capture different resolutions. For each level, we tested a range of 2 factors around it and used the number that led to the highest modularity. We used the scikit-learn Python package.
- **scHPF**: We ran scHPF on raw counts. Similarly to NMF, we used 30, 20, 10, and 5 factors but instead of choosing the number at each level that led to the highest modularity, we took the one with the largest ELBO value. We used the implementation in the schpf Python package.
- **Scanorama**: We ran Scanorama with default parameters from the implementation in the scanpy Python package using the PCA representation of the log-normalized counts.
- **Harmony**: We ran Harmony with default parameters from the implementation in the harmonypy Python package.
- **scVI**: We ran scVI with default parameters from the implementation in the scvi-tools Python package, using the raw counts as input.
- **Differential expression**: For each Leiden clustering result (including after batch integration), we obtained gene signatures for each cluster by obtaining differentially expressed genes between each cluster and all the others using the Wilcoxon rank-sum test as implemented in the scanpy Python package.
- **Hierarchies from each method**: For all methods except scDEF, we obtain 4 clusterings at different levels of resolution without any explicit connection between the levels. To create hierarchies from them, we assign each cluster from layer *l* to a cluster at layer *l* + 1 based on the fraction of cells from the lower-level cluster that is present in each cluster at layer *l* + 1.

### Performance metrics

- **Adjusted rand index**: We compute the adjusted rand index using cluster assignments at the highest resolution level for each clustering method. For scDEF, scHPF, and NMF, cluster assignments are obtained by assigning cells to the factors they attribute the largest weights to.
- **Signature accuracy**: For each method, we first assign annotated cell groups to clusters according to their F1 scores. With these correspondences, we evaluate the quality of the gene signatures by computing the average overlap index between the gene signatures obtained for each cluster with ground truth marker gene lists. The overlap index is computed as the number of genes in a true marker gene list that appear in the gene signature, divided by the size of the true marker gene list.
- **Hierarchy accuracy**: For each method, we first assign annotated cell groups to clusters according to their F1 scores. For each level in both true and inferred hierarchies, we list all groups that are below the groups at that level, regardless of their intermediate hierarchical relationships. We compute the average Jaccard index score across all levels in the true hierarchy.
- **Hierarchical signature consistency**: For each learned hierarchy, we create baseline expected gene signatures by computing a mixture of the gene signatures of the groups in the next lower level in the hierarchy, weighted by the size of the cell groups that contain those signatures. We create these signatures with the signatures obtained by the methods, and compare them to the gene signatures they obtained at higher levels in their hierarchies by their average Jaccard indices.

### Simulated data

Current methods to simulate scRNA-seq data are unable to generate data in a hierarchical structure. We therefore follow the procedure described in [15], with modifications that allow us to simulate technical and biological batch effects. The procedure is illustrated in Supplementary Figure S7.

The simulated hierarchical structure is encoded by a tree that contains cell groups at three levels of resolution, with two, four, and eight cell groups (Figure S7 A). We first simulate scRNA-seq data for 1000 cells and 1000 genes for two groups, corresponding to the top layer in the hierarchy. We then simulated 500 genes for the second level in the hierarchy, for the same number of cells, this time generating four cell groups. We repeated this procedure for another 250 genes to simulate the final eight groups. We aligned the cells generated at each step in such a way that the hierarchical structure is preserved (Figure S7 B), removing cells in groups that were inconsistent with the specified hierarchy. This recipe results in a matrix of 1000 cells and 1750 genes.

We instruct Splatter to generate four experimental batches in order to produce gene expression data with technical batch effects. We do this only for the last step in the hierarchical data generation procedure, in such a way that all eight cell groups are present in all batches. In this multi-batch scenario, we generate 1000 cells per batch. In order to simulate distinct biological conditions across batches, we sub-sample the complete data matrix by keeping only four out of eight of cell groups present in all batches, with the other four only appearing in each batch randomly. This captures the challenging but common setting in which each batch corresponds to a different biological condition with its own set of cell populations.

To ensure that the simulations are realistic, we estimate the global Splatter parameters using a public data set from 10x Genomics containing 3000 peripheral blood mononuclear cells from a healthy donor [25].

We obtain ground truth gene signatures for each high-resolution group by performing a Wilcoxon rank-sum test on the simulated data without batch effects and keeping the top 10 differentially expressed genes in each group.

## Data availability

The 3k PBMCs data used in this study are available from 10X Genomics [25] and were obtained from the SeuratData R package [43].

The *Schmidtea mediterranea* data used in this study are described in [26] and were downloaded from https://shiny.mdc-berlin.de/psca/.

The two batches of healthy PBMCs data used in this study are described in the Broad Institute PBMC Systematic Comparative Analysis study [27], and were obtained from the SeuratData R package [43].

The PBMCs data from two lupus patients used in this study are described in [28], and were obtained from the SeuratData R package [43].

The HGSOC data used in this study are described in [29], and were downloaded from the Gene Expression Omnibus (GEO) with accession code GSE165897.

## Code availability

scDEF is freely available as a Python package in https://github.com/cbg-ethz/scDEF. The code repository also contains Jupyter notebooks to reproduce the real data analyses and a Snakemake [44] workflow for the simulation study shown in this paper.

## Competing interests

The authors declare that they have no competing interests.

## Supplementary material

**Supplementary Figure S1:**
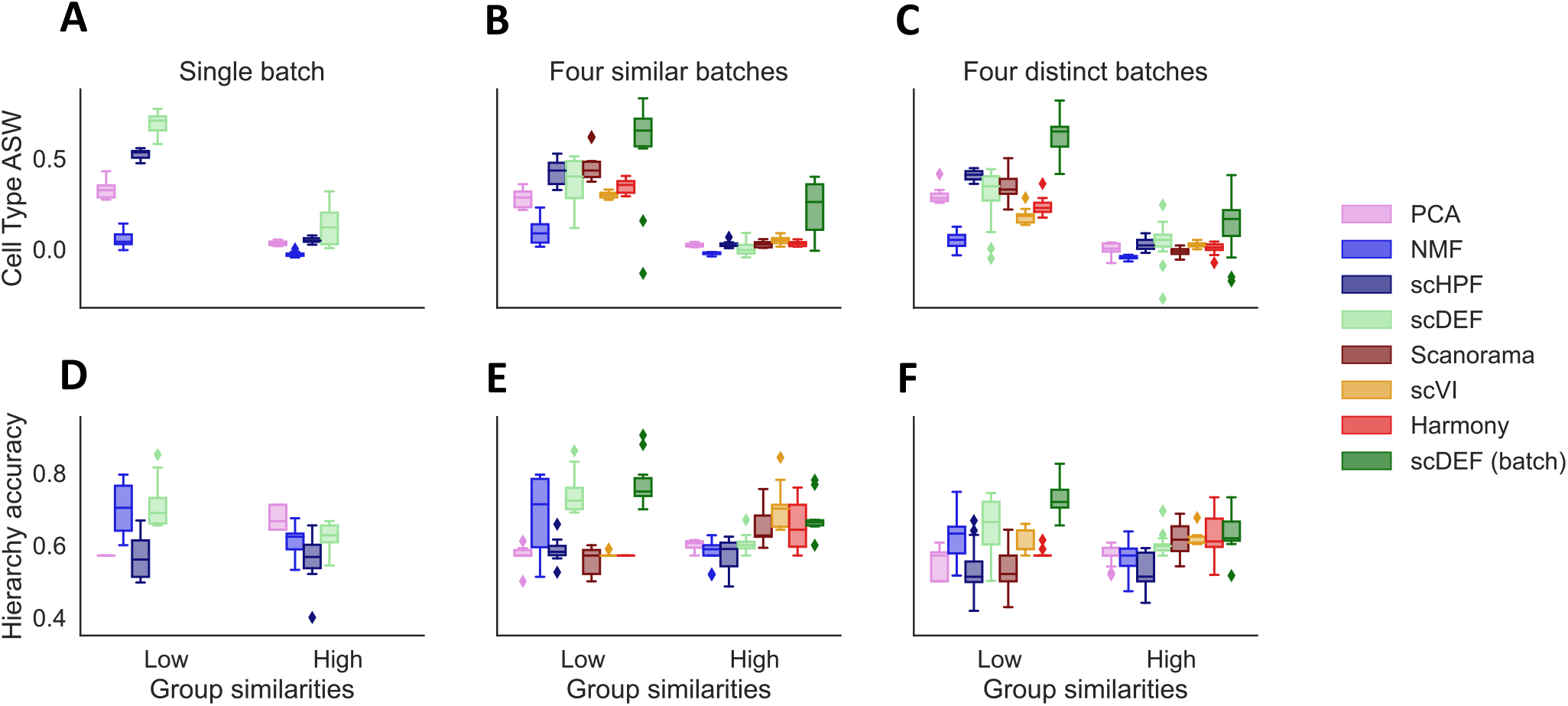
Average silhouette width (ASW) of the latent spaces learned by different methods and the accuracy of the hierarchies found by each one in simulated data.

**Supplementary Figure S2:**
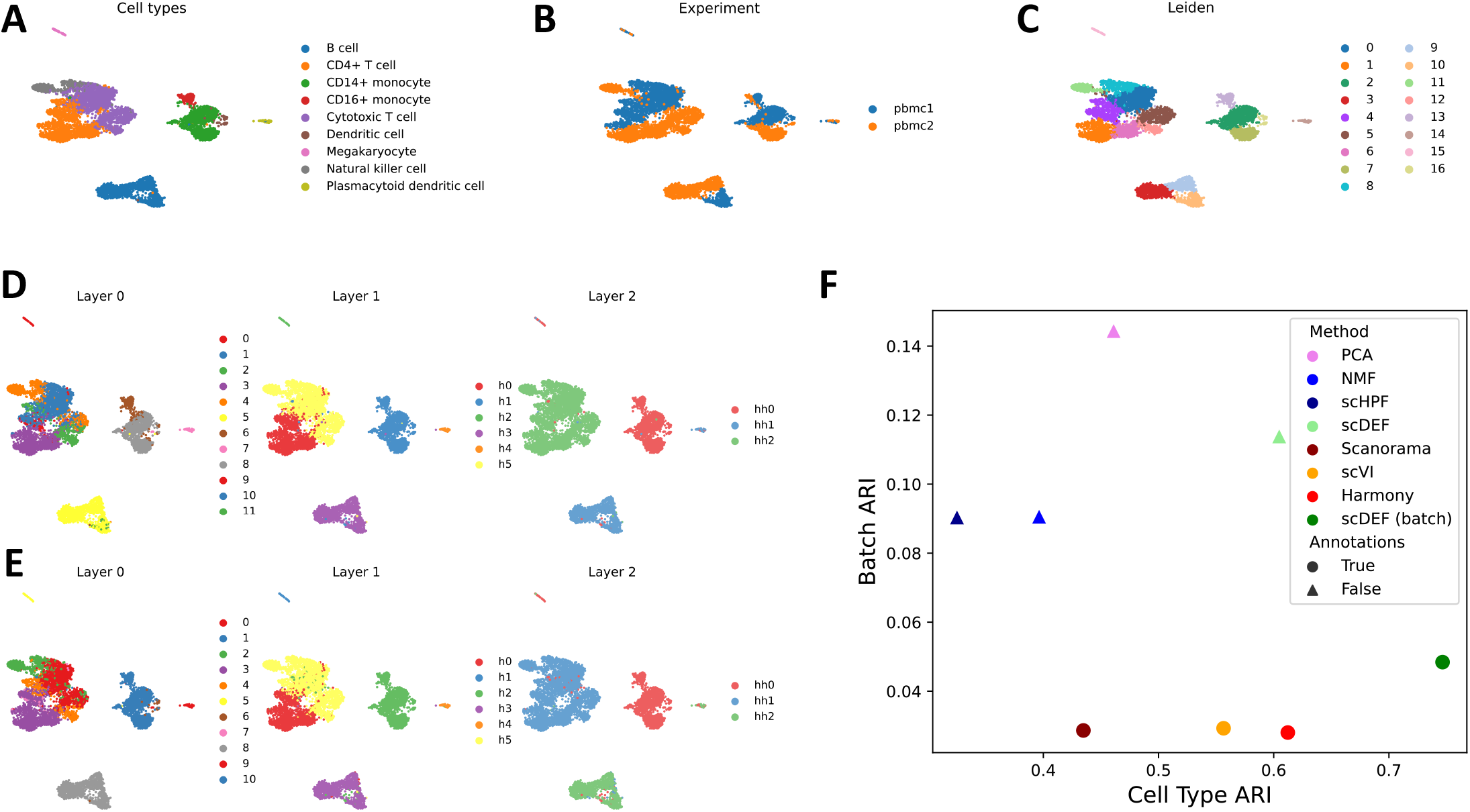
Application to 2 batches of healthy PBMCs. (A-E) UMAP embeddings of cells without any batch integration, coloured by (A) true cell type annotations, (B) batch of origin, (C) results from Leiden clustering without integration, (D) scDEF factors when batch annotations are not used, and (E) scDEF factors when batch annotations are used. (F) Comparison with other methods. ARI: adjusted rand index.

**Supplementary Figure S3:**
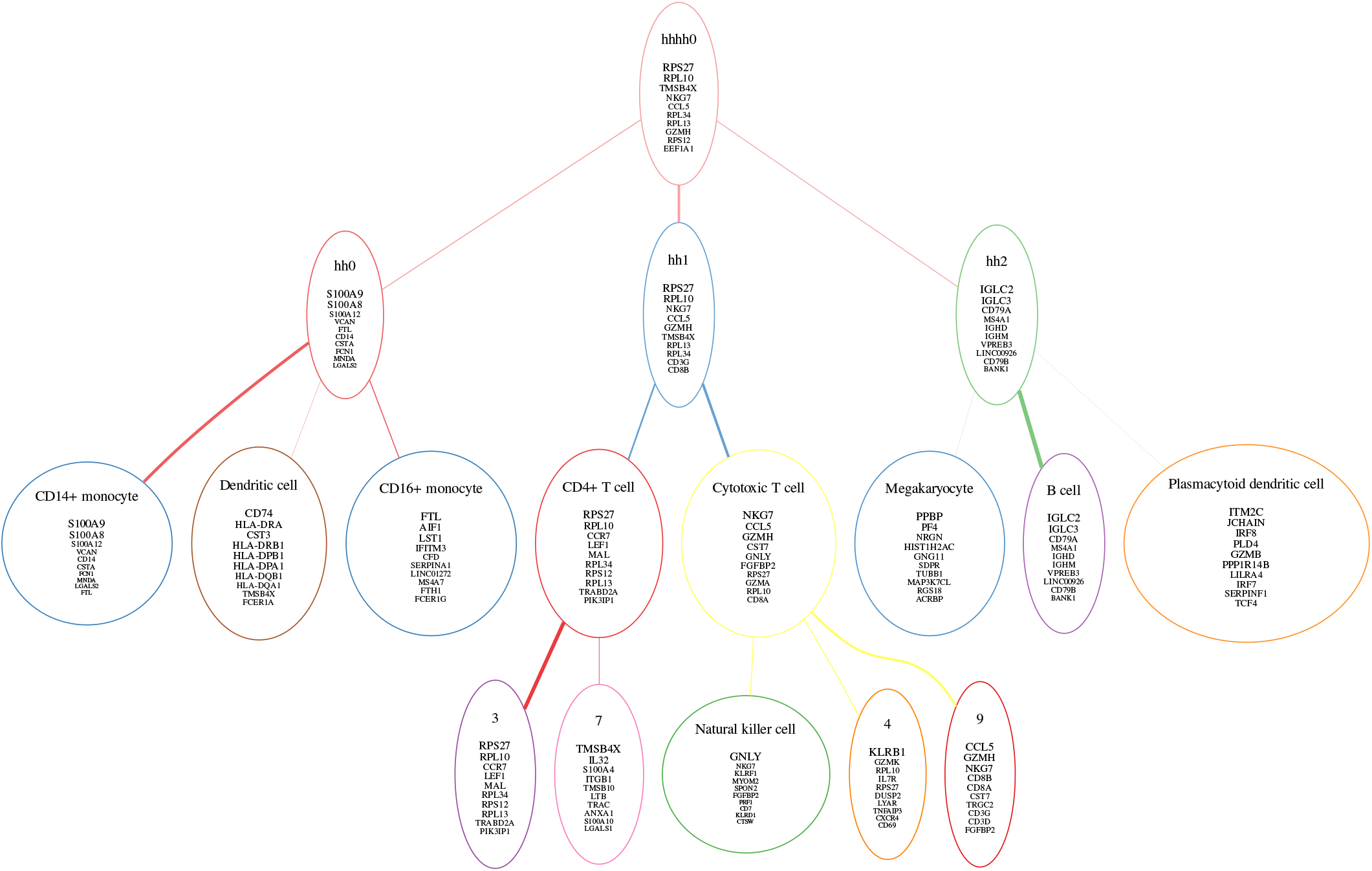
Application to 2 batches of healthy PBMCs. scDEF tree containing top 10 genes in each factor. Some factors are labeled by the provided annotated cell types that attach to each one. The node borders are coloured by the colors that correspond to each factor.

**Supplementary Figure S4:**
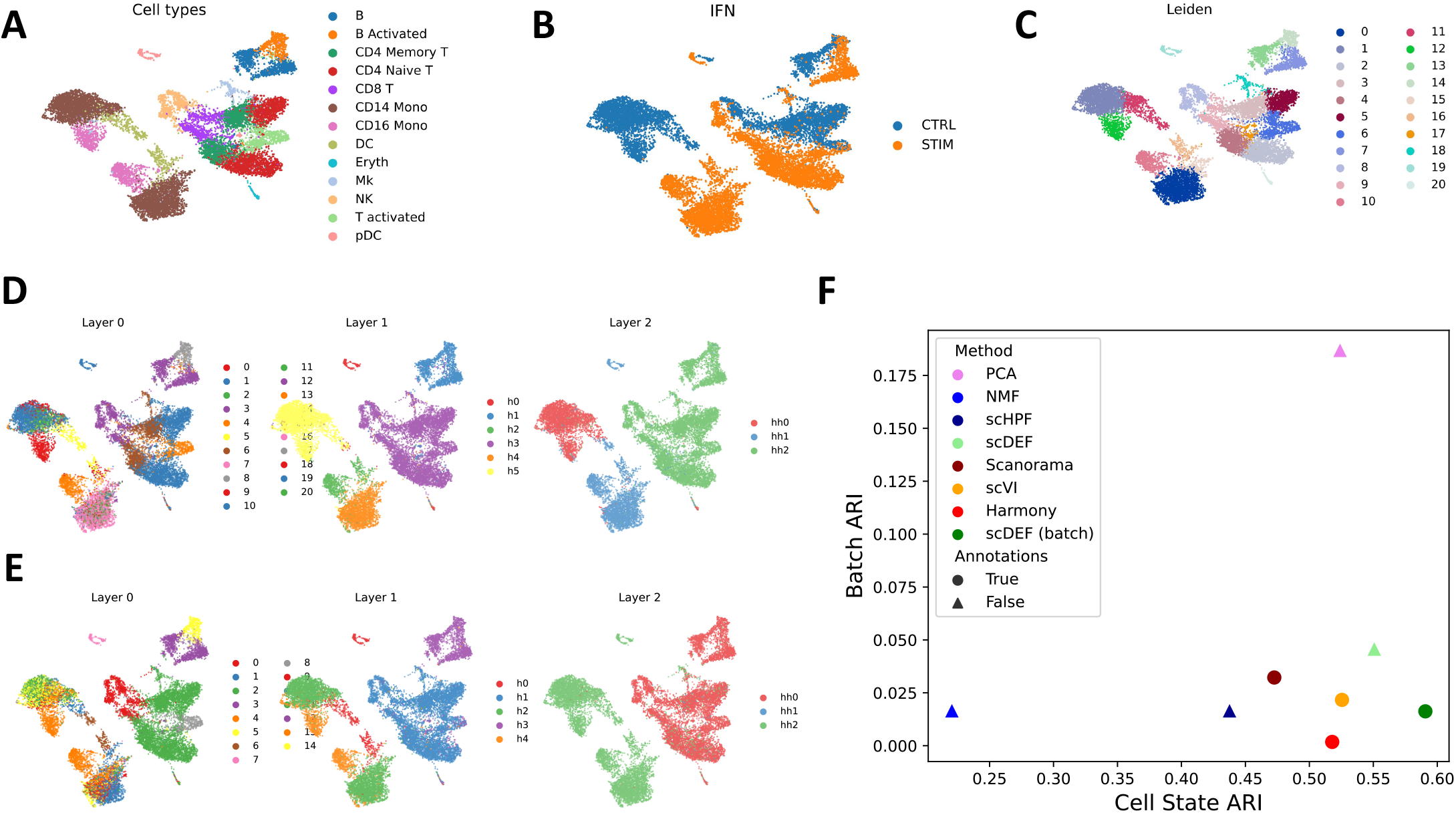
Application to 2 batches of healthy and interferon-stimulated PBMCs. (A-E) UMAP embeddings of cells without any batch integration, coloured by (A) true cell type annotations, (B) batch of origin, (C) results from Leiden clustering without integration, (D) scDEF factors when batch annotations are not used, and (E) scDEF factors when batch annotations are used. (F) Comparison with other methods. ARI: adjusted rand index.

**Supplementary Figure S5:**
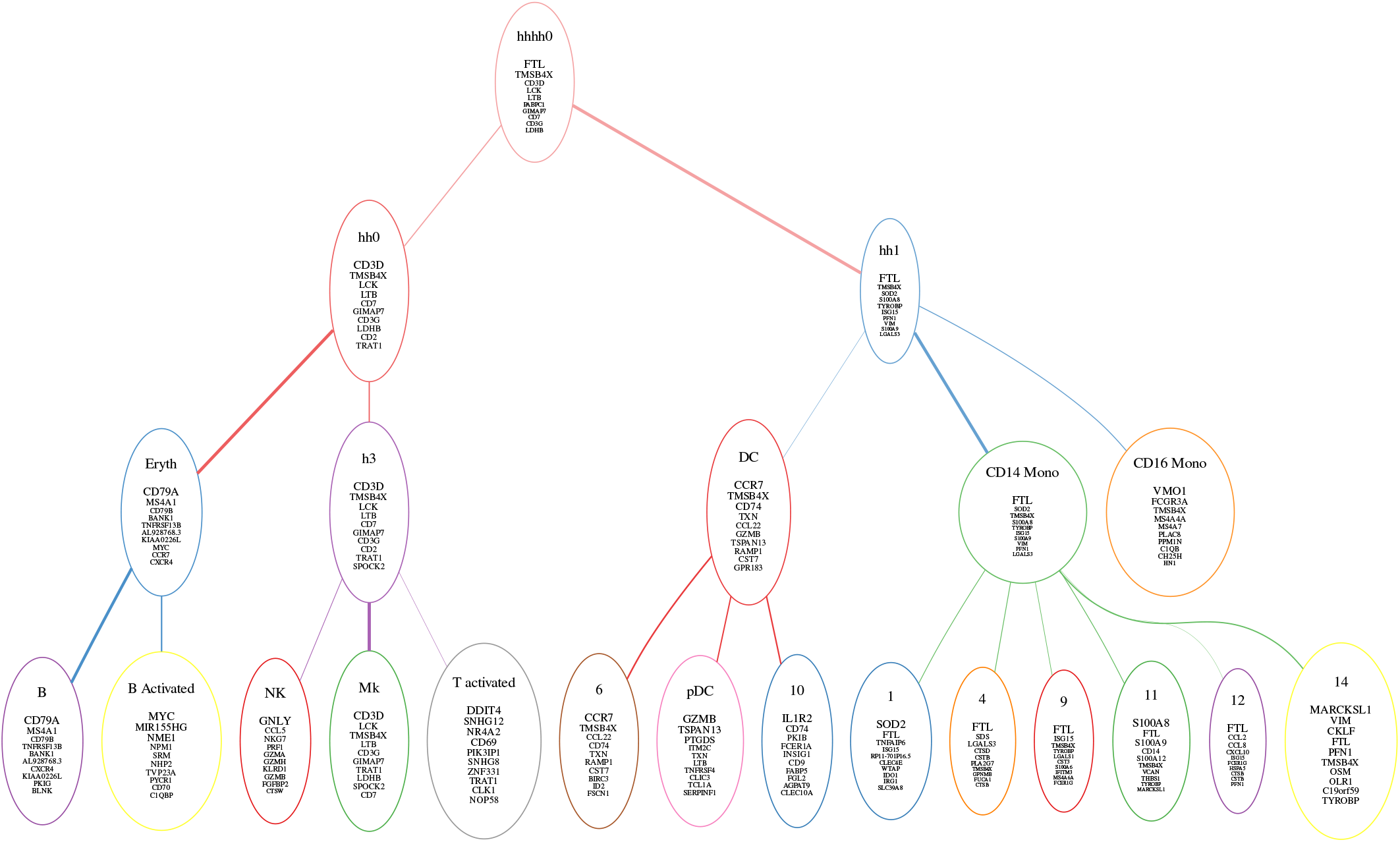
Application to 2 batches of healthy and interferon-stimulated PBMCs. scDEF tree containing top 10 genes in each factor. Some factors are labeled by the provided annotated cell types that attach to each one. The node borders are coloured by the colors that correspond to each factor.

**Supplementary Figure S6:**
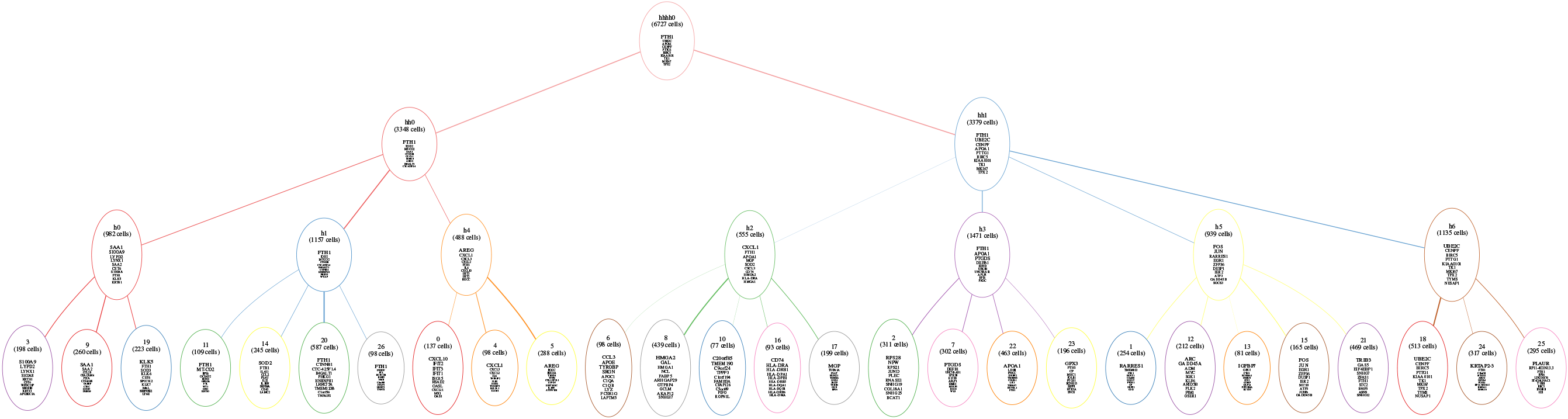
scDEF hierarchy obtained from 11 high-grade serous ovarian cancer patients with the top 10 genes in each factor displayed.

**Supplementary Figure S7:**
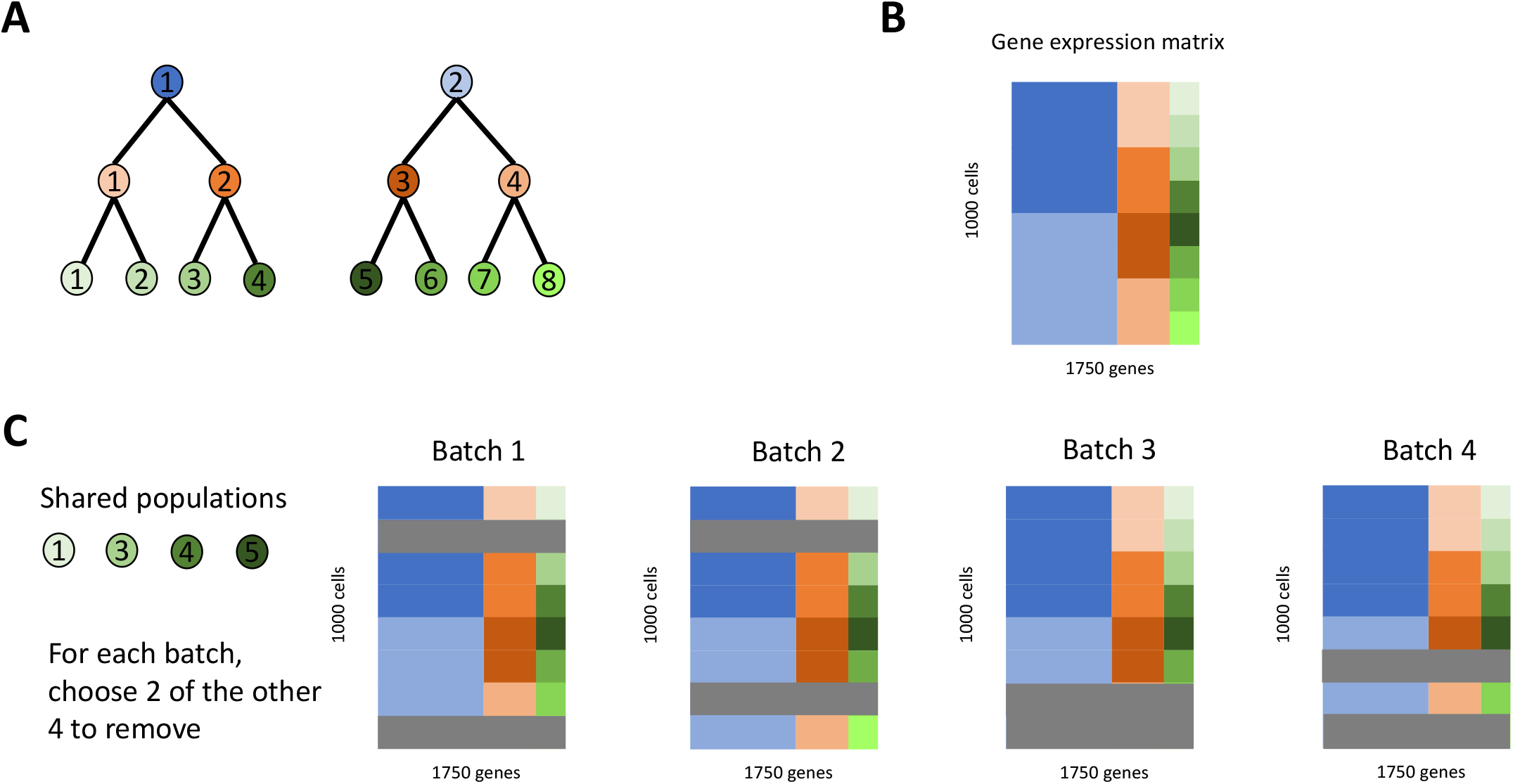
Generation of simulated data. (A) Hierarchical structure relating eight cell populations at three levels of resolution. (B) Simulated gene expression matrix containing 1000 cells and 1750 genes, where the first 1000 genes distinguish the top 2 groups, the next 500 distinguish the middle 4 groups, and the last 250 distinguish the bottom 8 groups. (C) For each generated data set in the distinct batch scenario, we randomly choose four cell groups in the highest resolution to be shared across all batches, and for each batch we randomly remove (in gray) two out of the remaining four.

### S1 Analysis of scRNA-seq data from 3k PBMCs

We applied scDEF to the publicly available 3k PBMCs data set from 10x Genomics, which contains 2623 peripheral blood mononuclear cells from a healthy donor [25]. This data set is often used for benchmarking scRNA-seq workflows as it contains well-defined cell types for which marker genes have been derived [18, 45]. We downloaded the processed data using Seurat [18], which contains the cell type annotations obtained from the canonical workflow of dimensionality reduction, clustering, and differential gene expression analysis. PBMC populations have a natural hierarchy consisting of three main groups containing T cells, B cells and monocytes, each of which may have their own cell subtypes or states with known marker genes, making this an appropriate data set to illustrate scDEF.

We ran scDEF with default parameters (Methods) and obtained a hierarchy containing three levels with 10, 5, and 4 factors for layers 0, 1, and 2, respectively, which enabled data visualization in 2D using UMAP embeddings and hierarchical signature discovery in an unsupervised manner (Figure S8). The Biological Relevance Determination prior (BRD, see Methods) guides the factorization towards solutions with factors that have a sparse profile, which we assume capture biological signal rather than technical variation (Figure S8 C). Indeed, taking the Gini index as a metric for sparsity (with higher values indicating that only a small number of genes are active in each signature), we find a strong correlation between the BRD posterior means and the Gini indices of the gene weights for each high-resolution factor (Figure S8 D). In this case, only 10 factors out of 100 from the lowest layer (Layer 0) were kept, and this selection is consistent with the selected factors from the upper layers due to the structure of the model.

The 2D projections of the cell weights at each layer indicate a gradual decrease in resolution as we progress from the lower to the upper layers of scDEF, with Layer 0 clearly capturing each annotated cell subtype, Layer 1 finding a factor corresponding to monocytes and another to cytotoxic T cells, and Layer 2 aggregating all T cells (Figure S8 B). The elongated embeddings of the cells using UMAP on scDEF factors contrast with the common compact profiles obtained by applying UMAP to PCA representations of the data. This is a natural consequence of the sparse scDEF factors, where each group of related cells will tend to attribute most of its weight to one factor, instead of a combination of factors. Furthermore, scDEF ensures that these latent representations are not influenced by library size and gene detection rate variations, which are due to technical and not biological effects, by estimating cell and gene-specific scale factors, which correlate strongly with the observed statistics (Figure S9).

By assigning each cell to the factor to which it attributes the most weight in each layer, we can inspect the correspondence between cell types and the scDEF factors (Methods, Figure S8 G). At the lower resolution levels, we founds that Layer 1 factor h2 has a high association score for both CD8 T cells and NK cells, factor h4 is associated with Naive and Memory CD4 T cells, and factor h0 captures monocytes. In the upmost layer, we obtain the strongest association score for T cells in factor hh1.

We next evaluated the quality of the gene signatures learned for each factor in the hierarchy (Figure S8 H). For each factor, we computed the overlap index between its gene signature and the marker genes of each annotated cell type (Methods). We found that the lower layer correctly identified marker genes for all populations, failing only at capturing the specific genes that are typically used to distinguish between Naive, Memory and CD8 T cells. This may be due to these genes being lowly-expressed even in these cells in this data set. As cell states get grouped together in the upper layers, the specific marker genes for some subtype with a lower abundance stop being captured in the top 20 genes, as expected. This is the case for FCGR3A+ monocytes in layer 1 and NK cells in layer 2.

The sparse nature of the weights that connect factors between adjacent layers in the scDEF model enables us to create a simplified tree structure (Figure S8 E) as follows: for each layer below the top one, we attach each factor to a factor in the layer right above it that it attributes the most weight to, and remove the other connections. We then also merge factors that contain only one child, as in, for example, factors 1, h3, and hh2, which all capture Platelets (Methods). This provides us with a compact representation of the hierarchical structure of the data.

scDEF is a Bayesian model which enables us to imbue the gene signatures in the hierarchy with confidence estimates (Methods). After associating factors with the annotated cell types, we may label each node in the hierarchy with its gene signature and confidence estimate. For example, the hierarchy shows factors corresponding to CD14+ Monocytes, FCGR3A+ Monocytes and Dendritic Cells, and their top 10 scoring genes, grouped together under a factor that contains a consensus signature of all the monocytes (Figure S8 F), as expected. The full annotated tree (Supplementary Figure S10) confirms that scDEF learned in an unsupervised fashion a biologically meaningful hierarchy of cell states in this data set.

#### S1.1 Incorporating biological information

While scDEF can be used in a purely unsupervised mode, often there are cell types with known marker genes that we wish to identify in our data, which is a common case for PBMCs, for example. We modify the prior distribution of the parameters of scDEF in such a way that they are guided towards the given fixed gene sets. To distinguish from the standard unsupervised model, we refer to this informed version as iscDEF (Figure S11). This version may use marker genes for highly resolved cell states and learn an upper level relation between them (Figure S11 A). Using canonical marker genes for the cell types provided with the annotations, scDEF correctly assigns cells to their true cell types and additionally aggregated them in a hierarchy that reflects their functional relationships accurately (Figure S11 B). Alternatively, iscDEF may use markers for coarse cell types and find more detailed cell states within them (Figure S11 C). In this case, we provided iscDEF with three gene sets corresponding to T cells, B cells and Monocytes, and instructed it to place them at the top level of the hierarchy. iscDEF then not only assigned cells to these coarse types but additionally learned cell-type specific states that strongly associated with the subtypes provided as ground truth annotations, resulting in a similar hierarchy to the one obtained in the fully unsupervised manner (Figure S11 D).

**Supplementary Figure S8:**
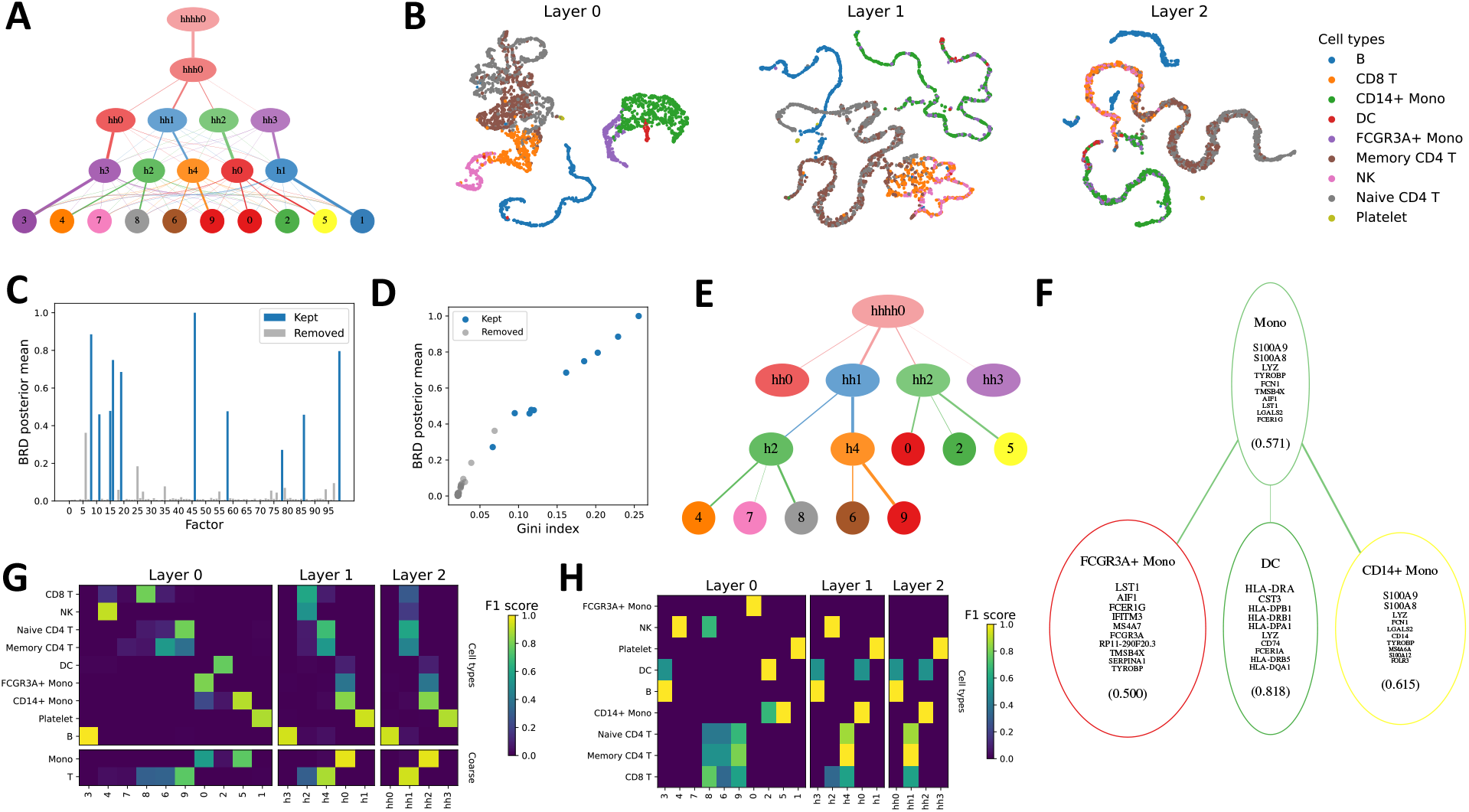
Application to 3k PBMCs. (A) The learned scDEF hierarchy. (B) UMAP embeddings of cells based on their weights at each scDEF layer, coloured by ground truth cell type annotation. (C) Posterior mean of the BRD variables. (D) Correlation between BRD posterior means and their Gini indices. (E) The tree obtained by assigning factors in each scDEF layer to the factors in the adjacent upper layer. (F) Factors strongly associated with the monocyte lineage show monocyte subtypes and their gene signatures with confidence estimates. (G) Association scores between scDEF factor assignments and cell type annotations. (H) Association scores between the top 20 genes in each factor and cell type markers.

**Supplementary Figure S9:**
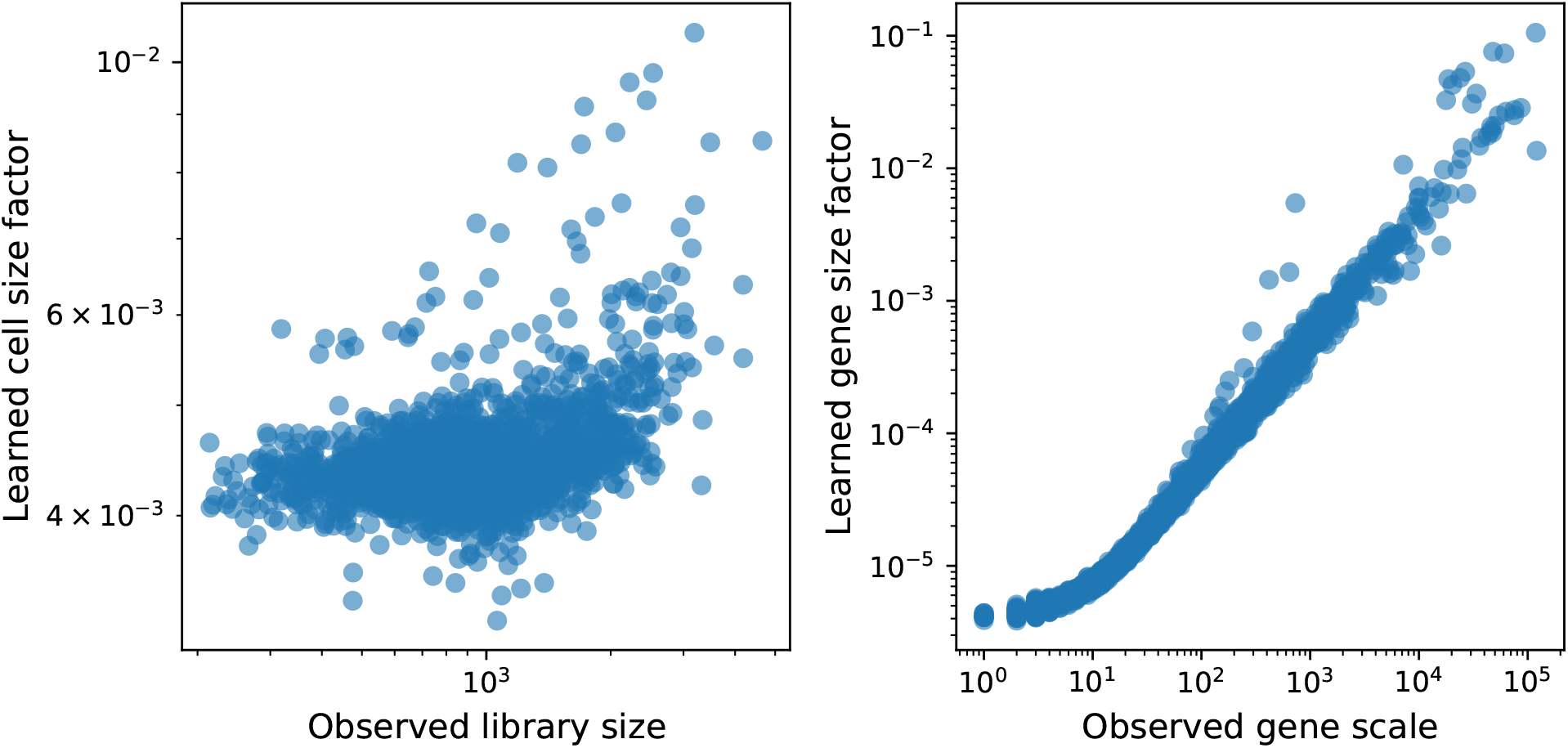
Application to 3k PBMCs. Comparison between the posterior means of the cell and gene scale parameters learned by scDEF and the observed library sizes and gene scales.

**Supplementary Figure S10:**
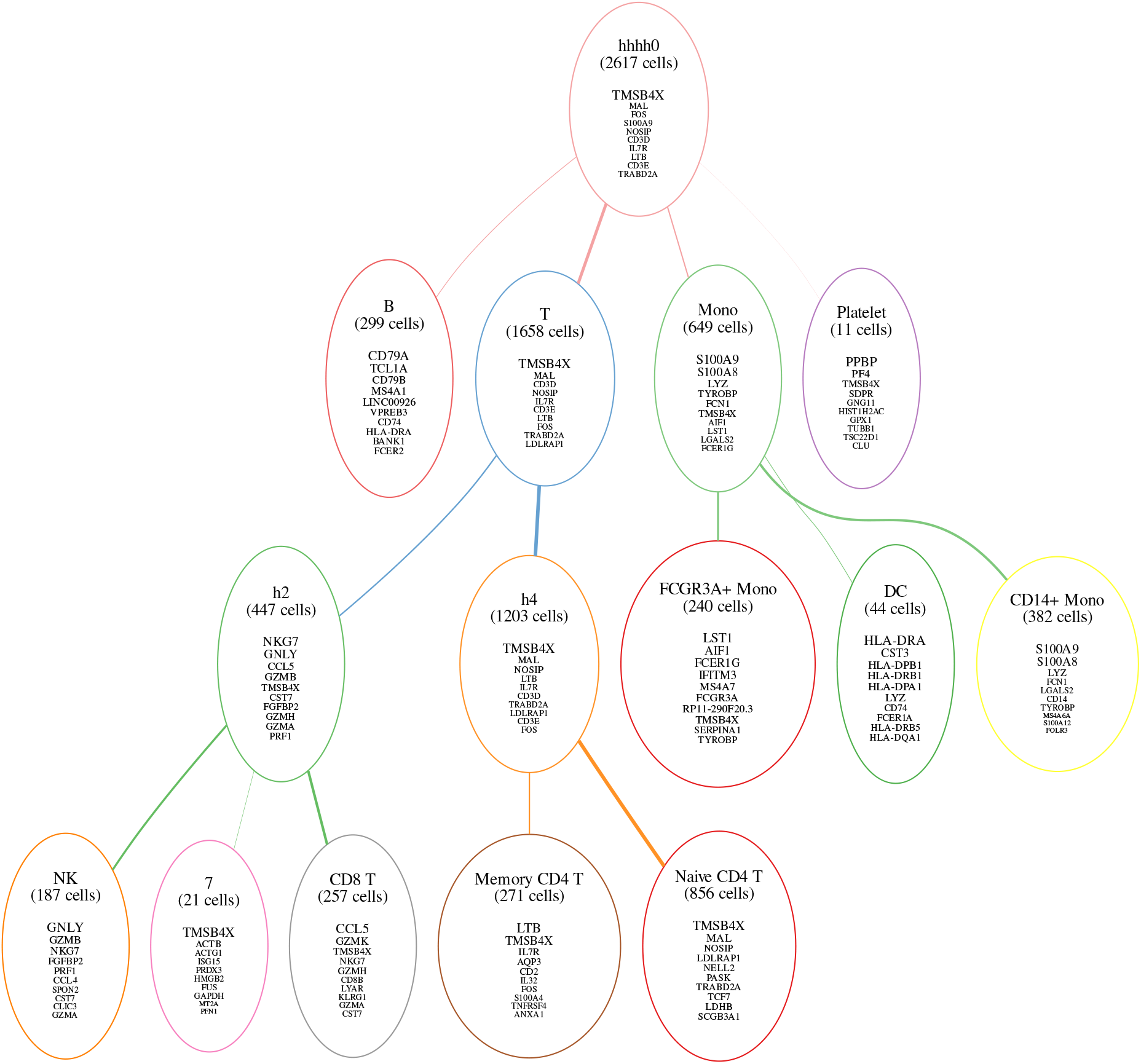
Application to 3k PBMCs. scDEF tree containing top 10 genes in each factor. Each factor is labeled by the provided annotated cell types that attach to each one, and by the number of cells it contains. The node borders are coloured by the colors that correspond to each factor.

This case study shows that scDEF can be used as a drop-in replacement to the usual scRNA-seq data analysis pipeline in a single-batch setting, enabling not only dimensionality reduction, unsupervised clustering, and gene signature identification, but also providing a meaningful hierarchy of cell states in the population we wish to study. Additionally, it provides in the same model the ability to use prior knowledge to guide the analysis, replacing the usual procedure in which users first assign cell types and perform clustering to identify higher-resolution states within coarse cell groups.

**Supplementary Figure S11:**
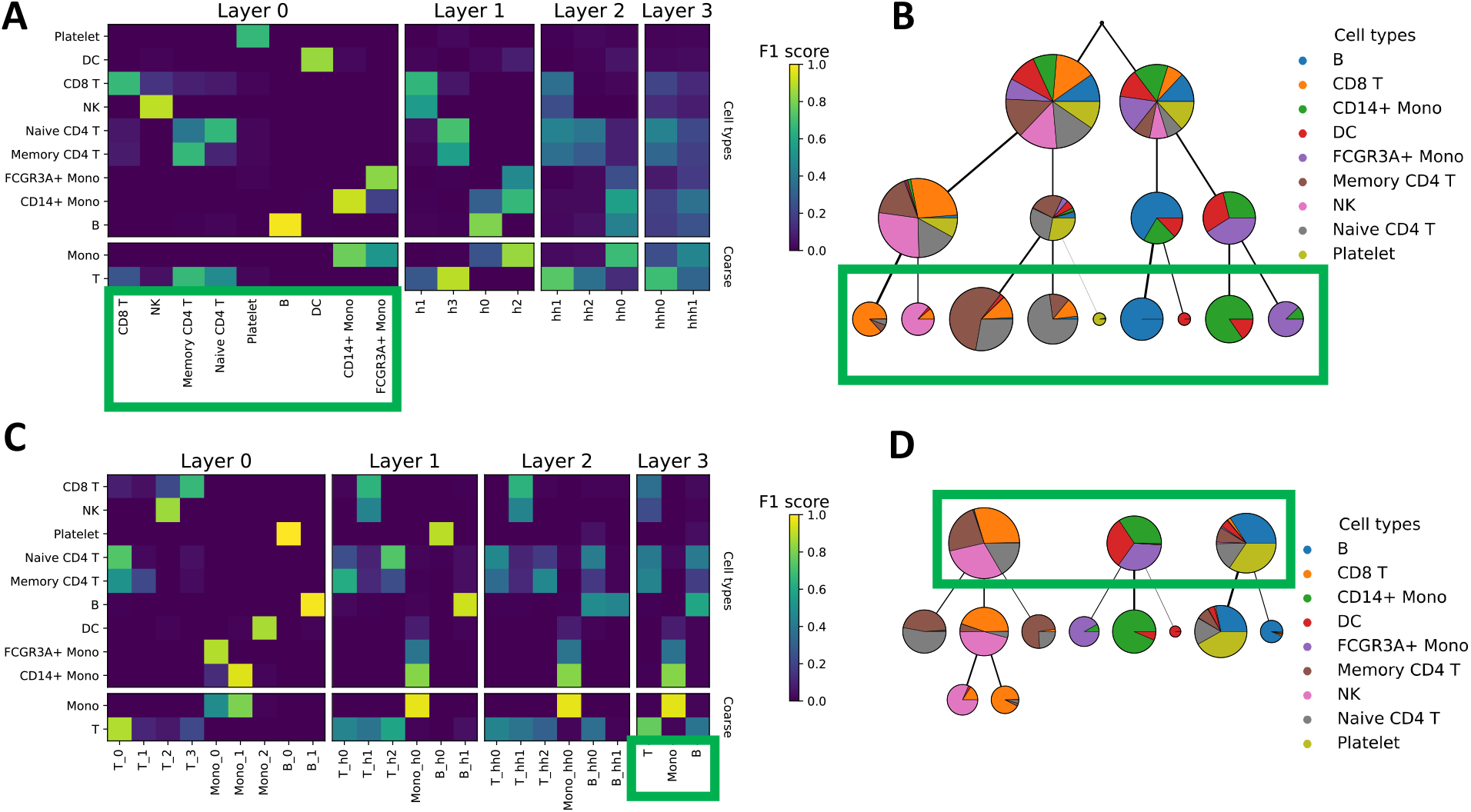
Application of informed scDEF (iscDEF) to 3k PBMCs. (A) Association scores between iscDEF factors and cell annotations, where iscDEF was guided by cell type markers at the lowest level layer and learned a hierarchy (B), where node sizes are scaled by the number of cells that attach to them and coloured by the proportions of each annotated cell type in the cells they contain. (C) Association scores between iscDEF factors and cell annotations, where iscDEF was used guided by cell type markers at the topmost level layer and learned higher resolution cell states (D) consistent with known biology. Green boxes indicate the hierarchy level at which iscDEF used gene sets as input.

## Notes

### Competing Interest Statement

The authors have declared no competing interest.

### Summary of Updates

Extended the model, added comparisons with more alternative methods, added more complete simulation study, and added more real data results.

