## Supplementary material for "Identifying hierarchical cell states and gene signatures with deep exponential families for single-cell transcriptomics"

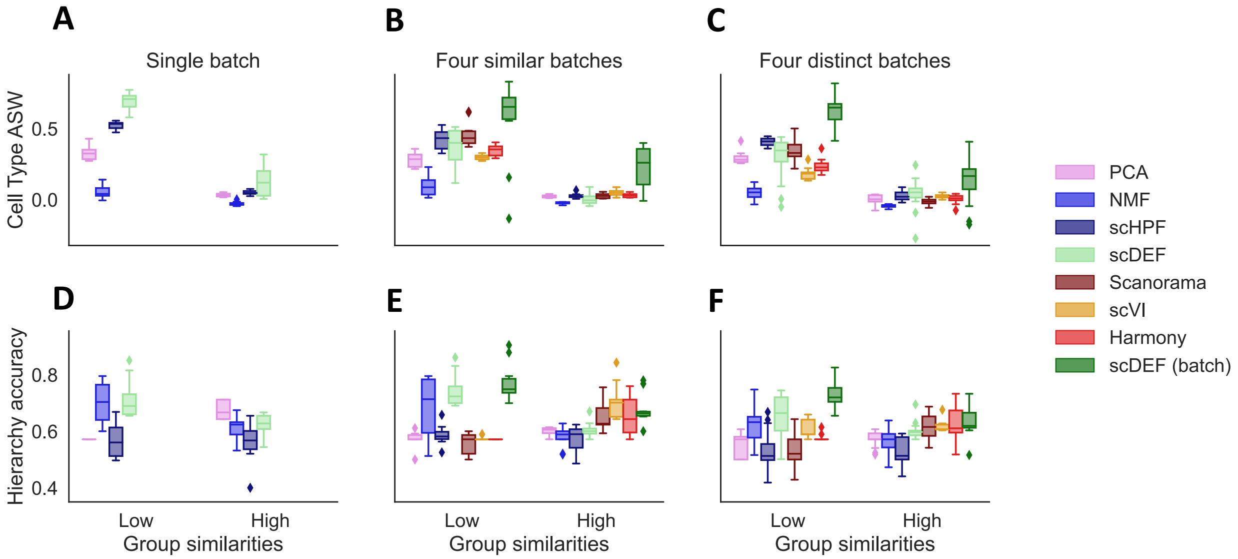


Supplementary Figure S1: Average silhouette width (ASW) of the latent spaces learned by different methods and the accuracy of the hierarchies found by each one in simulated data.

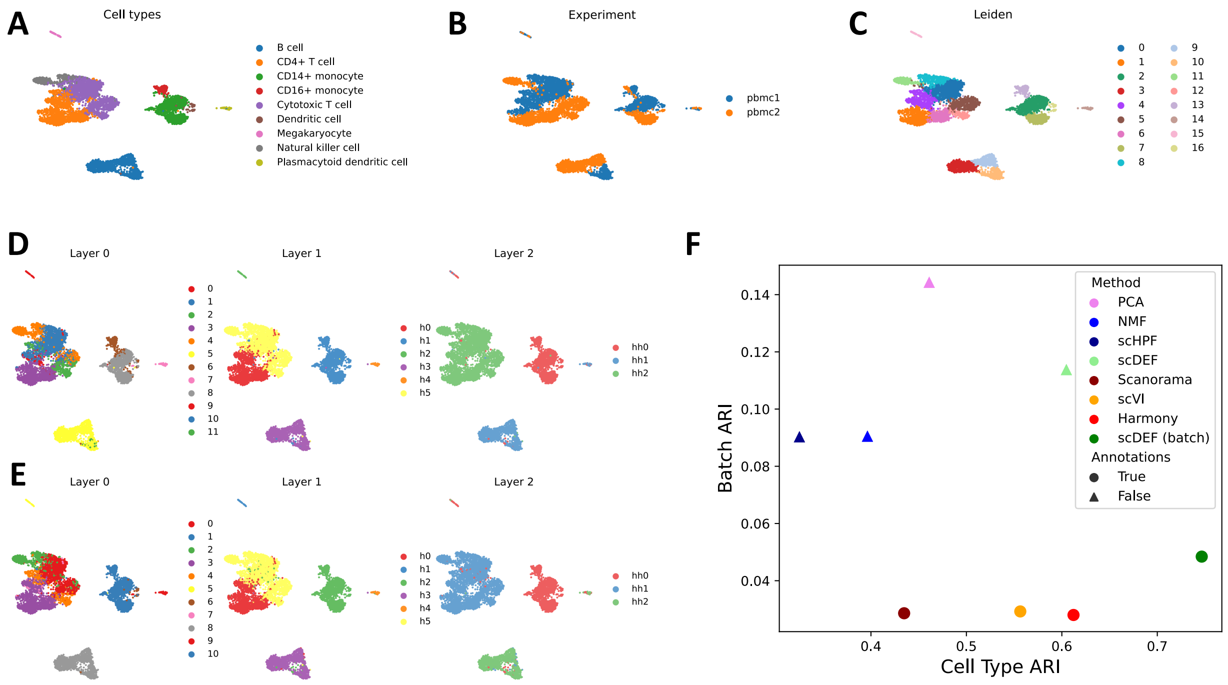


Supplementary Figure S2: Application to 2 batches of healthy PBMCs. (A-E) UMAP embeddings of cells without any batch integration, coloured by (A) true cell type annotations, (B) batch of origin, (C) results from Leiden clustering without integration, (D) scDEF factors when batch annotations are not used, and (E) scDEF factors when batch annotations are used. (F) Comparison with other methods. ARI: adjusted rand index.

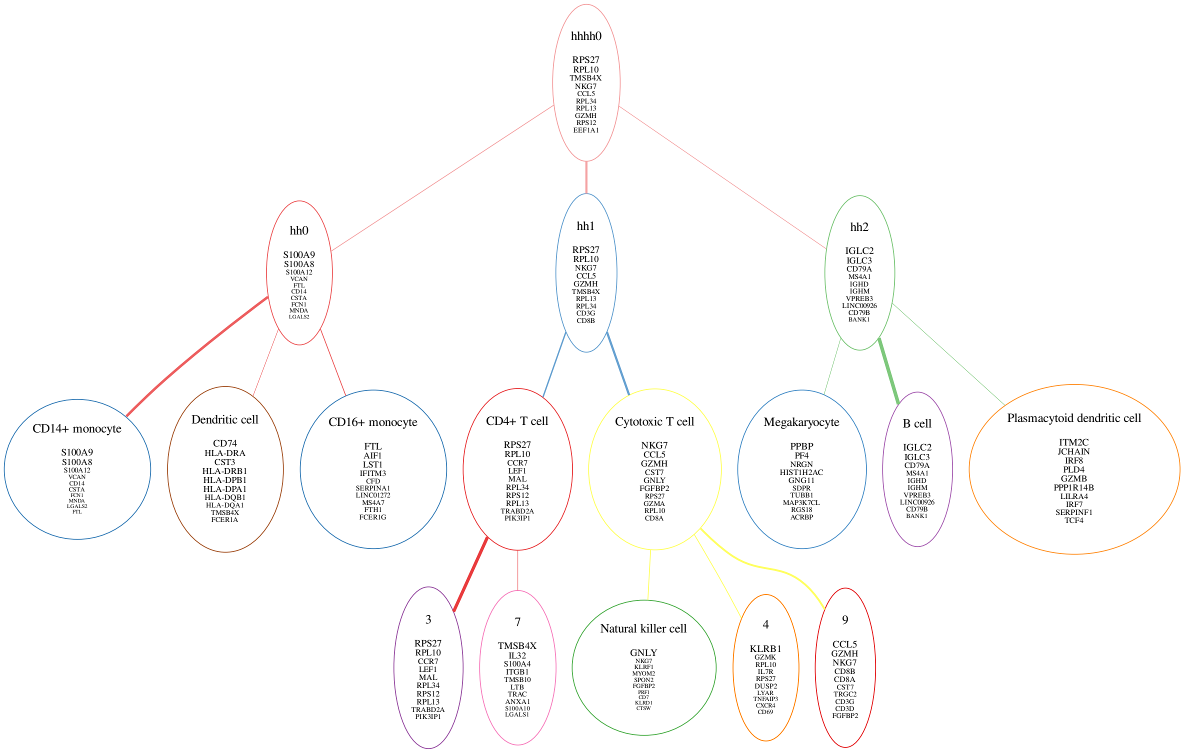


Supplementary Figure S3: Application to 2 batches of healthy PBMCs. scDEF tree containing top 10 genes in each factor. Some factors are labeled by the provided annotated cell types that attach to each one. The node borders are coloured by the colors that correspond to each factor.

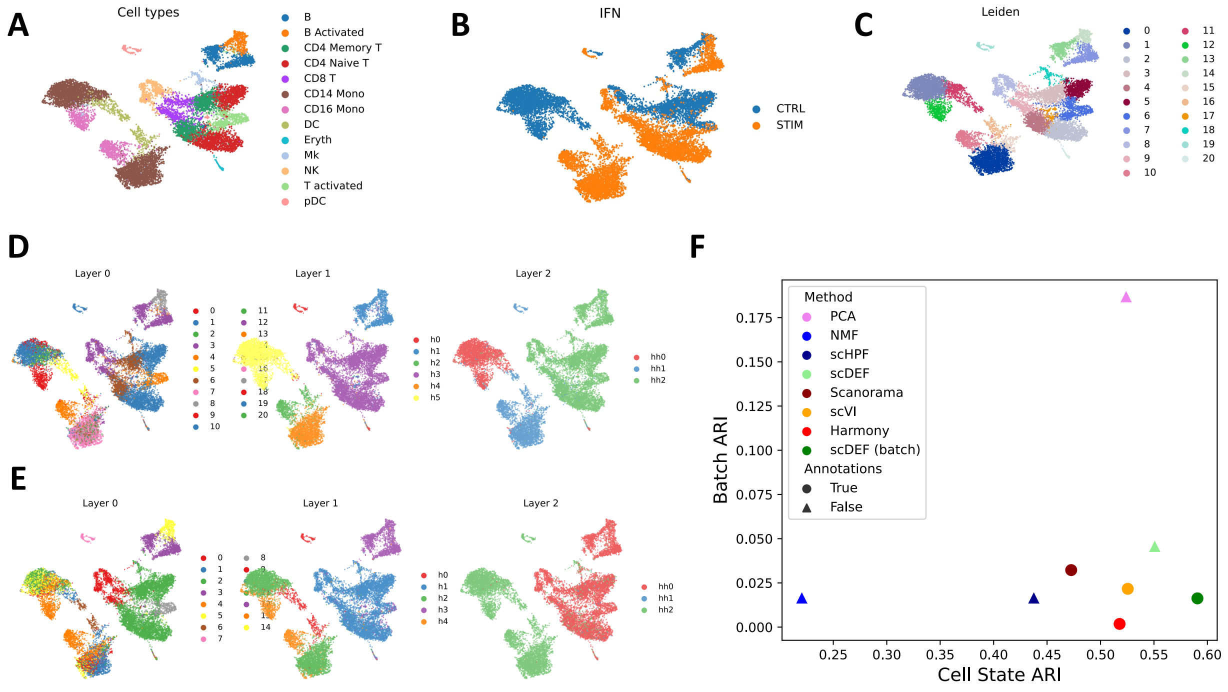


Supplementary Figure S4: Application to 2 batches of healthy and interferon-stimulated PBMCs. (A-E) UMAP embeddings of cells without any batch integration, coloured by (A) true cell type annotations, (B) batch of origin, (C) results from Leiden clustering without integration, (D) scDEF factors when batch annotations are not used, and (E) scDEF factors when batch annotations are used. (F) Comparison with other methods. ARI: adjusted rand index.

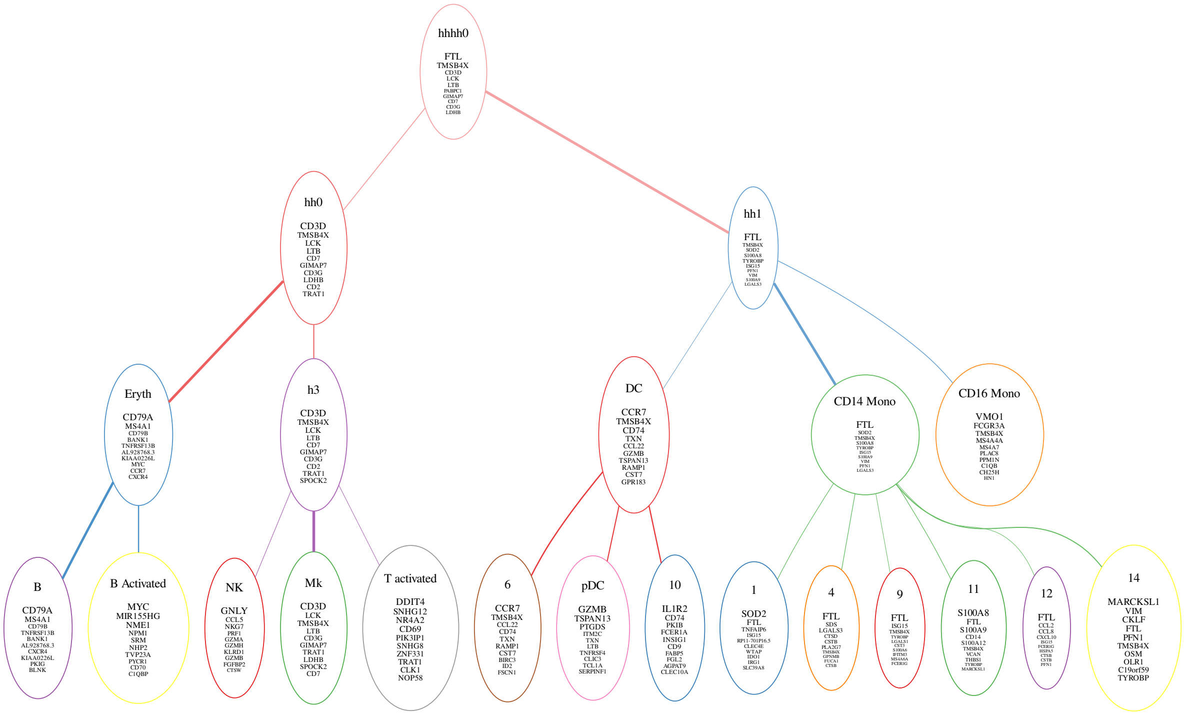


Supplementary Figure S5: Application to 2 batches of healthy and interferon-stimulated PBMCs. scDEF tree containing top 10 genes in each factor. Some factors are labeled by the provided annotated cell types that attach to each one. The node borders are coloured by the colors that correspond to each factor.

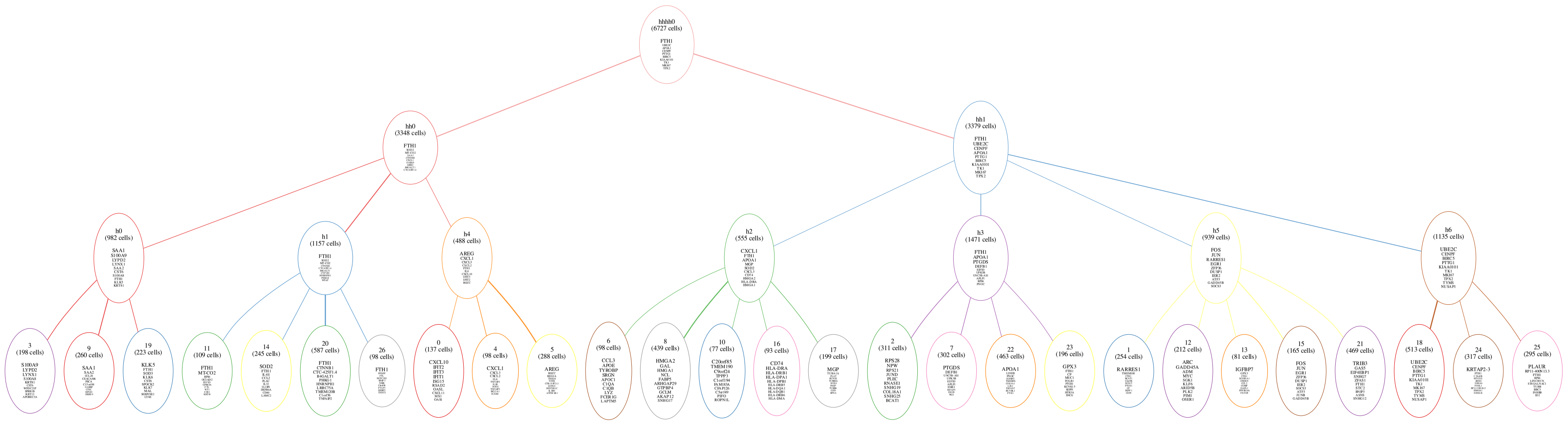


Supplementary Figure S6: scDEF hierarchy obtained from 11 high-grade serous ovarian cancer patients with the top 10 genes in each factor displayed.

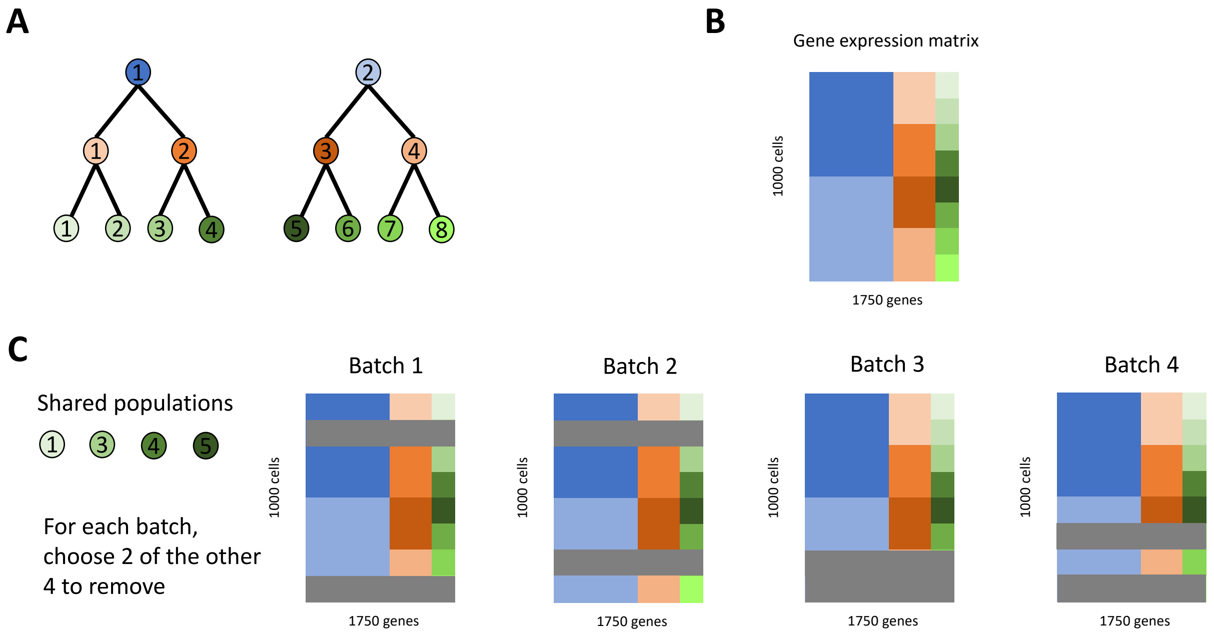


Supplementary Figure S7: Generation of simulated data. (A) Hierarchical structure relating eight cell populations at three levels of resolution. (B) Simulated gene expression matrix containing 1000 cells and 1750 genes, where the first 1000 genes distinguish the top 2 groups, the next 500 distinguish the middle 4 groups, and the last 250 distinguish the bottom 8 groups. (C) For each generated data set in the distinct batch scenario, we randomly choose four cell groups in the highest resolution to be shared across all batches, and for each batch we randomly remove (in gray) two out of the remaining four.

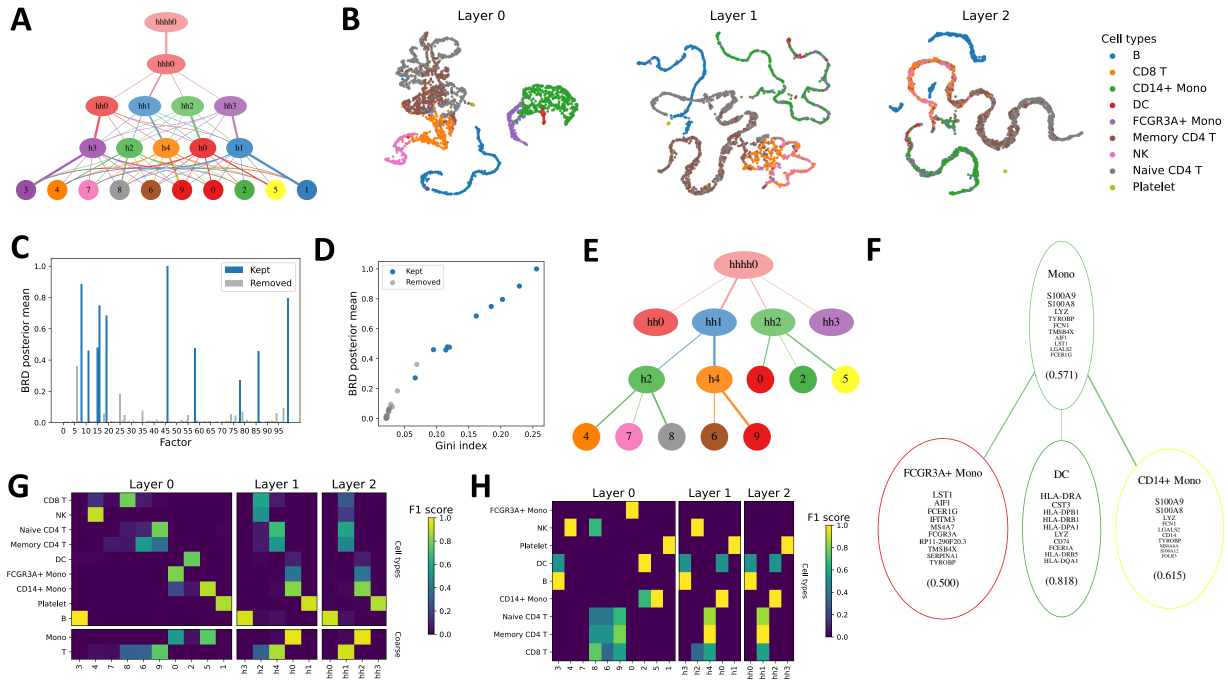


Supplementary Figure S8: Application to 3k PBMCs. (A) The learned scDEF hierarchy. (B) UMAP embeddings of cells based on their weights at each scDEF layer, coloured by ground truth cell type annotation. (C) Posterior mean of the BRD variables. (D) Correlation between BRD posterior means and their Gini indices. (E) The tree obtained by assigning factors in each scDEF layer to the factors in the adjacent upper layer. (F) Factors strongly associated with the monocyte lineage show monocyte subtypes and their gene signatures with confidence estimates. (G) Association scores between scDEF factor assignments and cell type annotations. (H) Association scores between the top 20 genes in each factor and cell type markers.

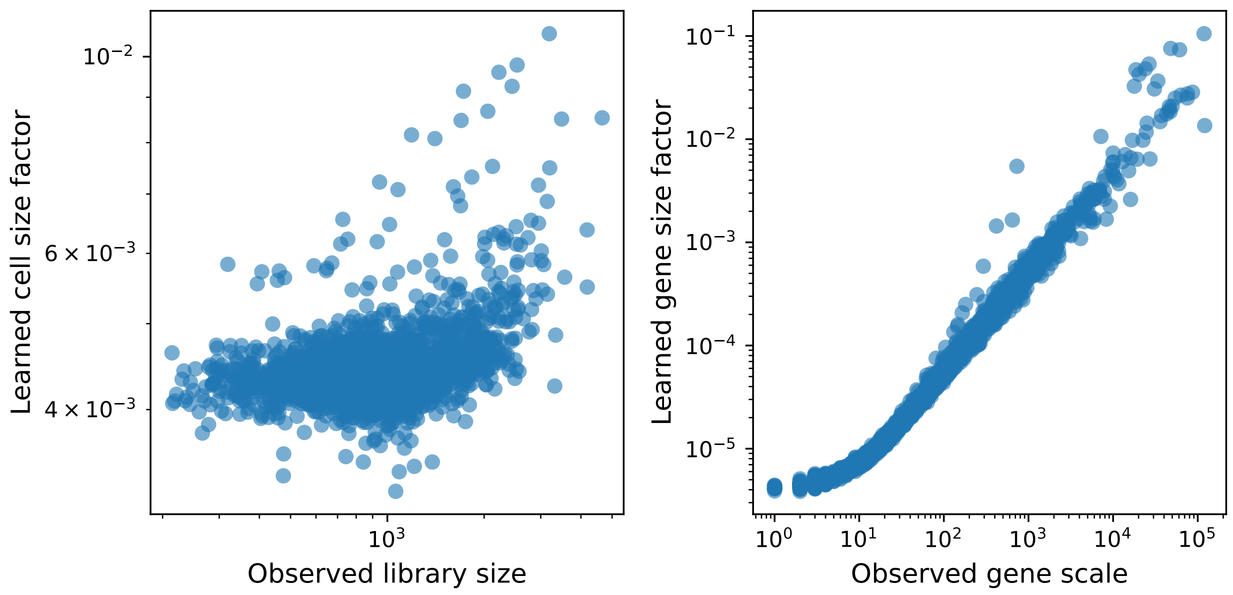


Supplementary Figure S9: Application to 3k PBMCs. Comparison between the posterior means of the cell and gene scale parameters learned by scDEF and the observed library sizes and gene scales.

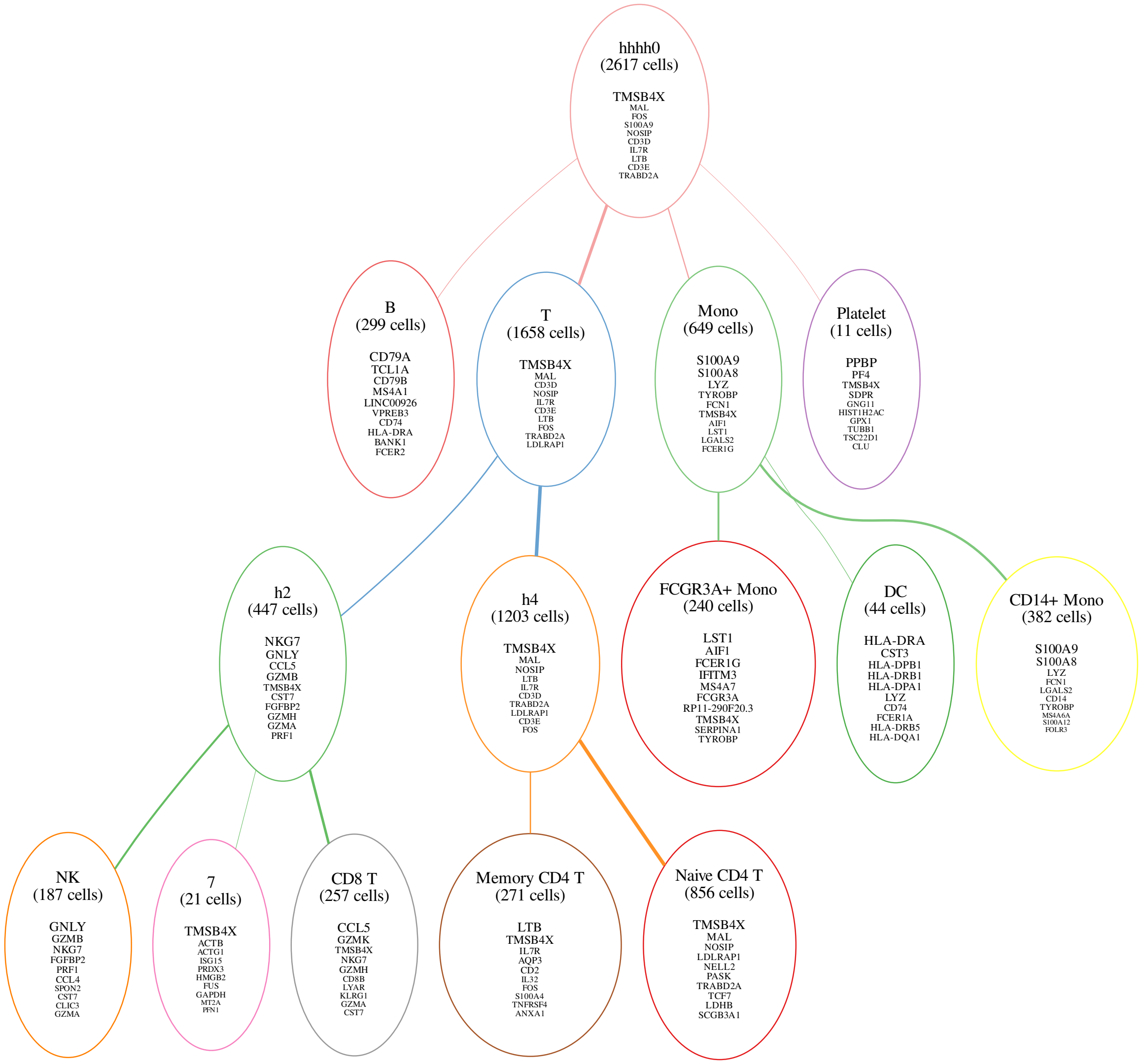


Supplementary Figure S10: Application to 3k PBMCs. scDEF tree containing top 10 genes in each factor. Each factor is labeled by the provided annotated cell types that attach to each one, and by the number of cells it contains. The node borders are coloured by the colors that correspond to each factor.

parameters of scDEF in such a way that they are guided towards the given fixed gene sets. To distinguish from the standard unsupervised model, we refer to this informed version as iscDEF (Figure S11). This version may use marker genes for highly resolved cell states and learn an upper level relation between them (Figure S11 A). Using canonical marker genes for the cell types provided with the annotations, scDEF correctly assigns cells to their true cell types and additionally aggregated them in a hierarchy that reflects their functional relationships accurately (Figure S11 B). Alternatively, iscDEF may use markers for coarse cell types and find more detailed cell states within them (Figure S11 C). In this case, we provided iscDEF with three gene sets corresponding to T cells, B cells and Monocytes, and instructed it to place them at the top level of the hierarchy. iscDEF then not only assigned cells to these coarse types but additionally learned cell-type specific states that strongly associated with the subtypes provided as ground truth annotations, resulting in a similar hierarchy to the one obtained in the fully unsupervised manner (Figure S11 D).

This case study shows that scDEF can be used as a drop-in replacement to the usual scRNA-seq data analysis pipeline in a single-batch setting, enabling not only dimensionality reduction, unsupervised clustering, and gene signature identification, but also providing a meaningful hierarchy of cell states in the population we wish to study. Additionally,

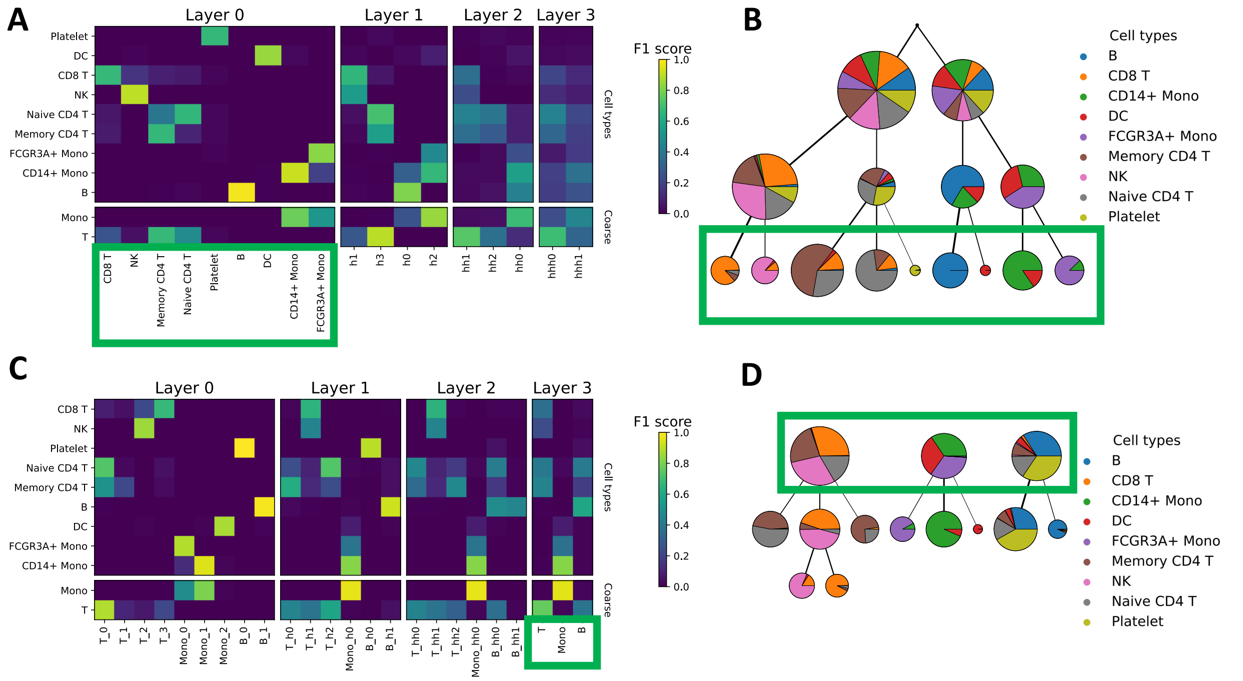


Supplementary Figure S11: Application of informed scDEF (iscDEF) to 3k PBMCs. (A) Association scores between iscDEF factors and cell annotations, where iscDEF was guided by cell type markers at the lowest level layer and learned a hierarchy (B), where node sizes are scaled by the number of cells that attach to them and coloured by the proportions of each annotated cell type in the cells they contain. (C) Association scores between iscDEF factors and cell annotations, where iscDEF was used guided by cell type markers at the topmost level layer and learned higher resolution cell states (D) consistent with known biology. Green boxes indicate the hierarchy level at which iscDEF used gene sets as input.

it provides in the same model the ability to use prior knowledge to guide the analysis, replacing the usual procedure in which users first assign cell types and perform clustering to identify higher-resolution states within coarse cell groups.
